# High potential contribution of intercropping to soybean and maize self-sufficiency in Europe

**DOI:** 10.1101/2024.12.17.627799

**Authors:** Mathilde Chen, Nicolas Guilpart, David Makowski

**Author notes:** Corresponding authors: Mathilde CHEN, PHIM, Univ Montpellier, CIRAD, INRAE, Institut Agro, IRD, 34398 Montpellier, France., David MAKOWSKI, Université Paris-Saclay, AgroParisTech, INRAE, UMR MIA PS, 91120 Palaiseau, France.

## Abstract

This study examines whether maize-soybean intercropping could improve the European Union’s soybean self-sufficiency while minimizing trade-offs with maize and other crops production. Assuming a four-year return, intercropping on 6.3, 13.1, and 20.1 Mha could produce 9.1 (28.8), 18.2 (63.7), and 27.2 (99.0) Mt of soybean (maize), achieving 25% (34%), 50% (75%), and 75% (116%) self-sufficiency. Reaching similar co-productions with sole crops require an additional 11.1, 24.7, and 36.5 Mha, implying 20-21% land savings with intercrops. Full soybean self-sufficiency could require more than 25 Mha of intercrops (25% of European cropland) annually. These estimates held under a large range of nitrogen inputs and temporal overlaps between crop species, provided 1 hectare of intercropping produced more soybean than 0.5 hectare of sole soybean. Geographic variation in sole crop yields should therefore be considered when assessing intercrops land-use efficiency. Our results highlight intercropping as a strategy to simultaneously improve European soybean production, satisfy maize needs, and save lands.

## Introduction

With the onset of multiple geopolitical crises, as well as volatility in food supply and prices,^1^ improving self-sufficiency has become a key priority of agricultural policies in the European Union (EU).^2,3^ Maize and soybean, two crops primarily used for livestock feed,^4,5^ are cultivated in the EU on 8.9 and 1.0 Mha, respectively (2018-2022 averages).^6^ However, domestic production quantities - 66.5 and 2.7 Mt -^6^ fall short of demand of 85.1 and 36.3 Mt, respectively.^7^ Although maize self-sufficiency is relatively high (81%), soybean self-sufficiency did not exceed 16% in 2021-2022.^8^ Moreover, EU’s imports of soybean are mainly sourced from a small number of countries. In 2023-2024, 98% of soybean imports came from Brazil (45.4%), the United-States (40.7%), Ukraine (7.4%), and Canada (4.0%).^9^ Increasing domestic soybean production is thus crucial to reduce EU’s dependency on massive imports and their negative environmental impacts in both exporting and importing countries.^10,11^

Previous studies suggest that satisfying 50% of the European consumption could require to grow 11.0-14.5 Mha of soybean,^12^ representing a substantial increase compared to the current soybean production area in Europe.^6^ Expanding croplands by converting natural ecosystems is unlikely, due to the scarcity of suitable land and the negative environmental impacts it could cause. Alternatively, soybean could substitute other crops, especially other spring crops with similar environmental niches such as maize.^12,13^ However, substituting soybean for maize could lead to a reduction of the total maize production in the EU. It is thus crucial to identify sustainable strategies that foster EU’s soybean self-sufficiency without compromising maize production.

So far, various scenarios of soybean area expansion in Europe have been explored in several studies, but they either overlooked the impact of soybean expansion on the production of other crops,^5,12,14,15^ or neglected intercropping,^13,16^ a practice still underdeveloped in Europe which consists in growing simultaneously multiple crops in the same field. Intercropping enhances land and nutrient use efficiency, making it a promising strategy for sustainable agricultural intensification.^17–19^ Based on production levels in China in 2018, estimates suggest that intercropping maize with soybean on 50% of the maize planting area could more than double the country’s total soybean production.^20^ An analysis of the potential of intercropping to sustainably boost the EU’s soybean self-sufficiency is still lacking.

In numerous field studies, the efficiency of intercropping over sole cropping is examined using the land equivalent ratio (LER), which corresponds to the sum of the ratios of intercrop to sole crop yields over the different species of the crop mixture. The ratio obtained for a given species, called the partial LER (pLER), represents the sole crop area that could be needed to obtain the same production as the intercrop.^21^ A global meta-analysis of 88 field experiments estimated that 0.56 ha of soybean and 0.79 ha maize grown separately (i.e., a total of 1.35 ha) were required to produce as much as 1 ha of maize-soybean intercropping,^22^ suggesting high potential of intercropping for land-saving compared to sole cropping. However, these field-scale estimates do not explicitly represent the spatial variability in maize^23^ and soybean^12–14^ yields across areas of contrasted productivity (e.g., due to climate conditions). Compared to intercropping, single cropping systems offer greater flexibility in terms of spatial distribution, as they allow production to be optimized by assigning each crop to its most productive area. If high-yielding areas in Europe are partly distinct for two crops, single cropping could allow farmers to allocate each crop to its highest-yielding region, while intercropping could involve growing at least one crop in low-yielding areas. It is therefore uncertain whether the large-scale adoption of intercropping could result in higher total production and land-use efficiency compared with sole cropping (see the detailed analysis of the productive performance of intercropping in case of spatial variability of yield in Supplementary Note 1).

In this study, we examine the potential of maize-soybean intercropping to improve soybean self-sufficiency in the EU while maintaining maize production. First, we estimate the area requirements under intercropping and sole cropping to meet 25%, 50% 75%, or 100% soybean self-sufficiency in the EU. Assuming that this level of soybean self-sufficiency is achieved, we then compare the efficiency of both strategies in maximizing both soybean and maize coproduction at the scale of the EU and quantify the land savings allowed by the most efficient strategy. The sensitivity of our results to variations in intercropping’s productive performances, return frequency, and area of cultivation is also explored. Our findings suggest that maize-soybean intercropping could substantially contribute to EU’s soybean self-sufficiency without decreasing maize production, while improving land use efficiency compared to sole cropping.

## Results

### Yield projections in the EU

Using historical (1981-2016) data of crop grain yield, water management, and climate in major producing areas of soybean and maize (Methods and Supplementary Figure 1), we trained machine learning algorithms to predict rainfed yields of soybean and maize at 0.5° grid-cell resolution across the EU as a function of local climate conditions (2000-2023). The developed models show good predictive performances, as illustrated by their mean R² values of 0.93 for soybean and 0.91 for maize obtained from two cross-validation procedures assessing temporal and spatial extrapolation (Supplementary Table 1 and Supplementary Figure 2) and by the symmetry of distribution of their residuals (Supplementary Figure 3). For both models, yield predictions are strongly impacted by total precipitation and water management, followed by temperatures (Supplementary Figure 4). Higher predicted yields are observed at higher precipitations and temperatures across the growing season, while lower yields are projected in dryer conditions (Supplementary Figures 5-9). These results are consistent with current knowledge on crop physiology of maize and soybean.^24,25^

We examine the suitability of soybean production in the EU using grid-cell-wise mean annual rainfed yield projections over 2000-2023. The 25^th^, 50^th^ (i.e., median), and 75^th^ percentiles of soybean productivity across the EU are 1.0, 2.1, and 2.6 t.ha^-1^ (Figure 1a, top panel), reached over 103, 82, and 36 Mha, respectively (Figure 1a, bottom panel). The level of productivity of 1.0 t.ha^-1^ also corresponds to the minimum yield observed in major soybean-producing regions with climatic conditions comparable to those of the EU in our train dataset (i.e., Italy and US). Considering this threshold as a criterion for soybean suitability, our projections suggest that nearly all European cropland areas (i.e., 109 Mha,^26^ including arable lands and permanents crops)^27^ could be considered as suitable for soybean production, greatly exceeding the current production area of 1 Mha (Supplementary Table 2). Compared to previous studies,^12–14^ we obtain a larger share of soybean-suitable area in northeastern Europe, especially in Germany and Poland, but lower suitability in Spain. Dominant crops currently cultivated in soybean-suitable area are wheat, maize, and barley (Supplementary Table 3). Maize shares a similar ecological niche with soybean,^28^ suggesting that it is the most likely crop species to be replaced as soybean area expands.

**Figure 1.**
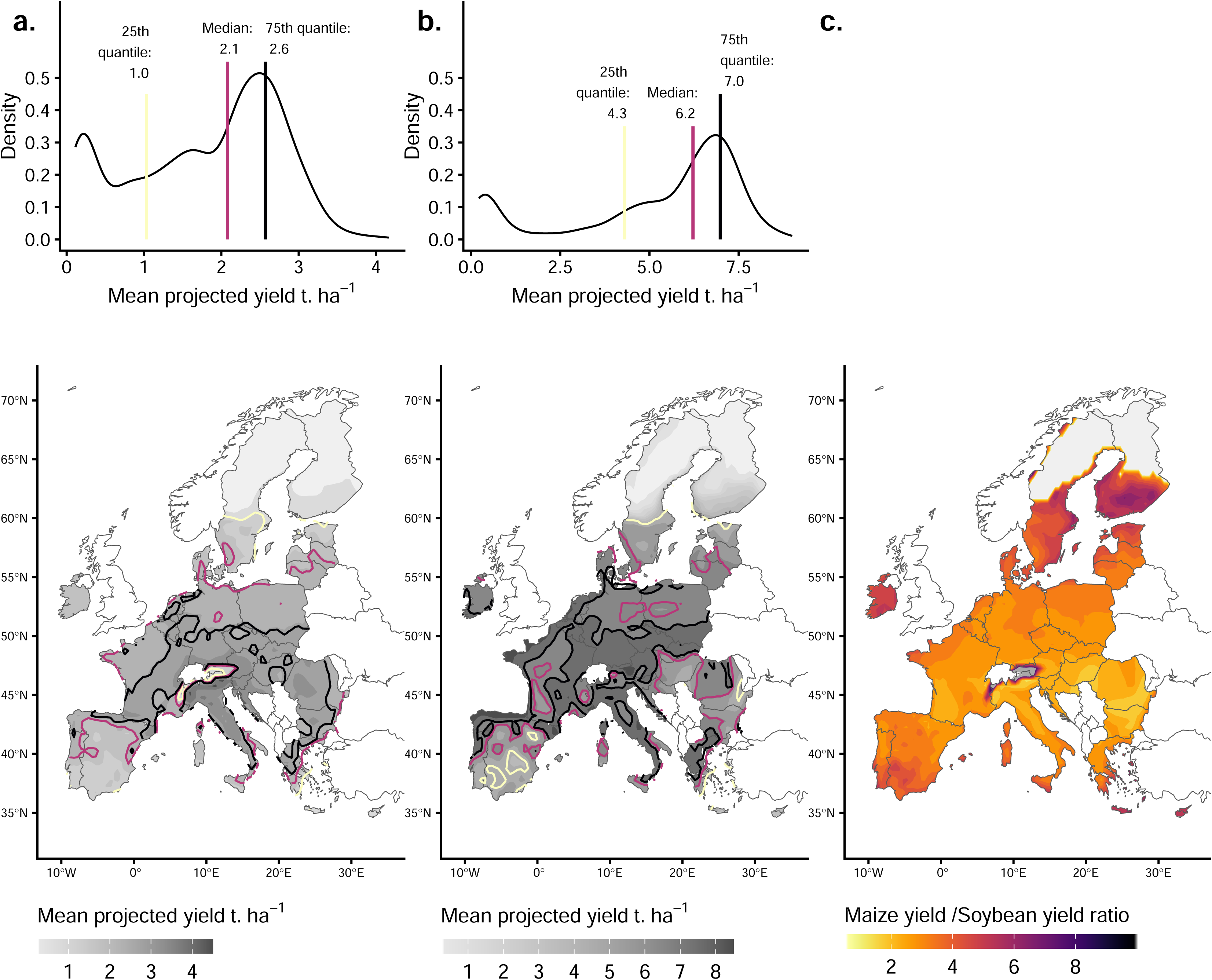
Soybean (a) and maize (b) rainfed yield projections (2000-2023 averages), and maize/soybean ratio (c) in the European Union. Projections were obtained from random forest models based on climate inputs (derived from the ERA5-land dataset) covering the crop growing seasons (April-November for soybean and April-December for maize) and irrigated fraction (estimated by SPAM2010, and set to zero for the yield projections). Climate inputs included monthly minimum and maximum temperatures, total precipitation, solar radiation, reference evapotranspiration, and vapor pressure deficit. For each climate variable, the first two scores derived from a principal component analysis were used as climate predictors in the model. For panels (a) and (b), density graphics on the top show the variability in average projected yields, with 25^th^, 50^th^ (median), and 75^th^ quantiles indicated as vertical yellow, red, and black lines, respectively. Below, the maps represent the spatial distribution of projected average yields, with darker shades indicating regions with higher mean yields, whereas lighter shades representing lower yields (e.g., areas with average crop yield ∼ 0 t.ha^-1^ are displayed in light grey). Regions corresponding to the 25^th^, 50^th^ (median), and 75^th^ yield quantiles are further delineated by yellow, red, and black contours, respectively. For panel (c), darker shades indicate high ratio between maize and soybean average projected yields, for example in the areas close to the Alps characterized by low soybean yields (i.e., <1 t.ha^-1^) but moderate-to-high maize yields (i.e., >6 t.ha^-1^). Base map based on Natural Earth data, created using the R package rnaturalearth.

For maize under rainfed conditions, the corresponding 25^th^, 50^th^, and 75^th^ percentiles of projected yields across the EU are 4.3, 6.2, and 6.9 t.ha^-1^ (Figure 1b, top panel), attained over areas of 100, 62, and 27 Mha, respectively (Figure 1b, bottom panel). Our projections are consistent with historical maize yields in Europe,^29^ except for Spain where our projections are slightly lower, maybe because of uncertainties about irrigation. High-yielding areas for both crops encompass a band in central EU stretching from southwestern France to the Balkan region, excluding the Alps (Figure 1c). On the contrary, areas with contrasting productivity between maize and soybean are found in Ireland, northern and southern EU.

### Land savings allowed by intercropping over sole cropping at a continental scale

Following the procedure displayed in Figure 2 and detailed in the Supplementary Note 2, we examine the potential of maize-soybean intercropping to improve soybean self-sufficiency in the EU while maintaining maize production.

**Figure 2.**
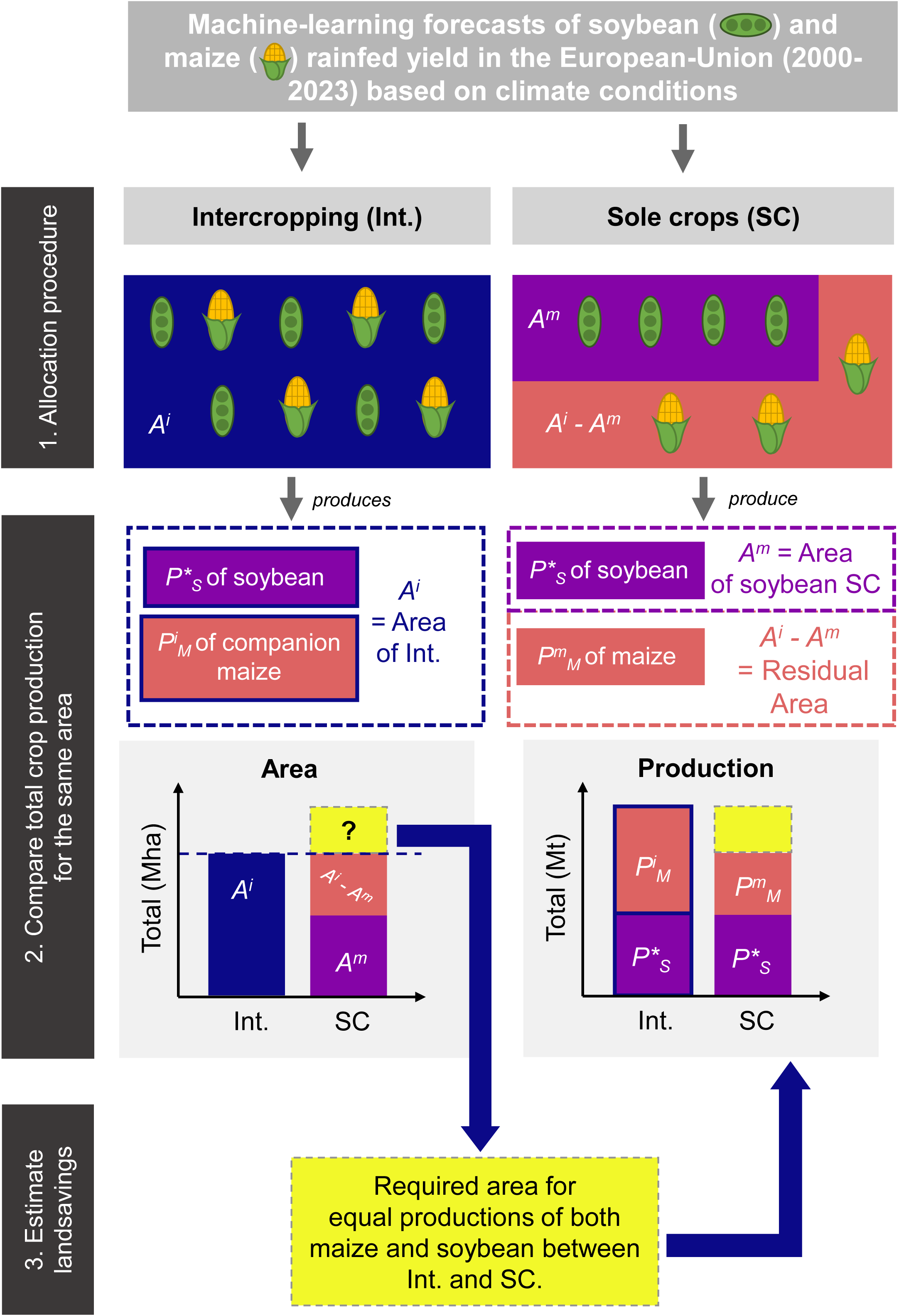
Procedure applied to estimate the area requirements under intercropping (Int) and sole cropping (SC) to meet a target soybean production P*_S_ in the European Union (EU), to compare the efficiency of both strategies in maximizing both soybean and maize coproduction, and to quantify the land saving allowed by the most efficient strategy at the scale of the EU. (1) Maize-soybean Int. and SC are sequentially allocated in areas ranked by soybean production, if climatic conditions are suitable for the production (≥1 t.ha^-1^) until P^∗^ is reached or if the total area allocated to intercropping (i.e., A^i^) exceeds 25 Mha. (2) The total soybean and maize productions from both strategies are compared (P*_S_ Mt of soybean and P^i^_M_ Mt of maize produced on A^i^ Mha for Int. vs P*_S_ Mt of soybean produced on A^m^ Mha and P^m^_M_ Mt of maize produced on A^i^-A^m^ Mha for SC). (3) Across the EU, the required area for equal maize and soybean productions between Int. and SC. is then estimated.

First, we combine our yield projections with the productive performances of intercropping to estimate the potential quantity of soybean produced across the EU by a reference maize-soybean intercropping system. To represent the current average productive performance of maize-soybean intercropping systems, we assume as a baseline that 1 ha of intercropping produces as much as 0.56 ha of soybean and 0.79 ha of maize grown separately,^22^ corresponding to pLER values of 0.56 and 0.79, respectively. We consider that intercrops are grown once every four years,^30,31^ which means that the area cultivated under maize-soybean intercropping in a given site each year cannot exceed 25% of its cropland area.

We then estimate the area requirements under such a system to achieve 25%, 50%, 75%, and 100% soybean self-sufficiency in the EU (i.e., 9.1, 18.2, 27.2, and 36.3 Mt, respectively), assuming that intercropping is primarily implemented in regions with the highest soybean productivity. We sequentially allocate intercropping to the most productive sites, ranked by soybean production (i.e. estimated by multiplying projected sole soybean yield by the cropland area in each site), until the whole intercropping area produces a quantity of soybean covering the considered level of self-sufficiency (i.e., 25%, 50%, 75%, or 100%) or exceeds 25 Mha, an area nearly equivalent to a quarter of European cropland areas.^32^ We define this limit to avoid any extreme land use scenario leading to strong reductions of areas allocated to other crops (Supplementary Table 3), and because it is compatible with a soybean frequency of one-in-four year. Since intercropping involves that both soybean and maize are cultivated together in the same field, we also estimate the quantity of maize produced in this area.

Table 1 presents the area requirements under maize-soybean intercropping to cover each targeted level of soybean self-sufficiency in the EU, as well as co-productions of both soybean and companion maize. For example, the total area allocated to maize-soybean intercropping to achieve 50% soybean self-sufficiency in the EU could cover 13.1 Mha, located mainly in southeastern part of France, North of Italy, Hungary, and Romania where the production of soybean is high (Supplementary Figure 10). This system could produce 18.2 Mt of soybean and 63.7 Mt of maize, representing nearly a sevenfold increase in current EU soybean production while maintaining maize production at nearly the same level (Supplementary Table 2). In other words, this system could cover 50% of the EU’s soybean needs and nearly 75% of its maize demand could be met, thereby more than tripling the current self-sufficiency rate for soybean (16%), while slightly reducing the current maize self-sufficiency rate (81%).

**Table 1.** Soybean and maize coproduction and area requirement for 25, 50, 75, and 100% soybean self-sufficiency in the European Union (EU) achieved by intercropping or sole cropping.

|  | Target % of EU's self-sufficiency in soybean <sup>a</sup> | Intercropping |  | Sole crops |  |  | Total |  | Land-saving <sup>b</sup> |
| --- | --- | --- | --- | --- | --- | --- | --- | --- | --- |
|  |  | Soybean | Maize | Soybean | Maize | Additional land required | Intercrop-ping | Sole crop |  |
| **Production (Mt)** | **25%** | 9.1 | 28.8 | 9.1 | 17.7 | 11.1 | 37.9 | 37.9 | NA |
|  | **50%** | 18.2 | 63.7 | 18.2 | 39.0 | 24.7 | 81.9 | 81.9 | NA |
|  | **75%** | 27.2 | 99.0 | 27.2 | 59.1 | 36.5 | 126.2 | 122.8 | NA |
|  | **100%** | 33.1* | 123.6 | 36.3 | 65.0 | 5.3 | 156.7 | 106.6 | NA |
| **Area (Mha)** | **25%** | 6.3 |  | 3.4 | 2.9 | 1.7 | 6.3 | 8.0 | 21% |
|  | **50%** | 13.1 |  | 7.1 | 6.0 | 3.4 | 13.1 | 16.5 | 21% |
|  | **75%** | 20.1 |  | 10.9 | 9.2 | 5.8 | 20.1 | 25.0 | 20%** |
|  | **100%** | 25.0 |  | 14.8 | 10.2 | 0.9 | 25.0 | 25.0 | 0%** |

Notes: Crop return frequency was one-in-four year for all simulations and total allocated area was limited to 25% of croplands in the EU (i.e., 25 Mha). For intercropping, partial land equivalent ratios were equal to 0.56 and 0.79 for soybean and maize, respectively.
<sup>a</sup> Self-sufficiency levels of 50, 75, and 100% correspond to 18.2, 27.2, 36.3 Mt soybean.
<sup>b</sup> Computed as the absolute relative difference of surface covered by sole crop vs intercropping: $\text{abs}[(\text{surface\_intercropping} - \text{surface\_sole cropping}) / \text{surface\_sole cropping}]$ .
\* Target production not reached even when soybean was grown of 25% cropland in the EU.
\*\* Production from intercropping strategy higher than sole crops.

Then, we estimate the area requirements to reach the same soybean self-sufficiency levels but under a sole cropping system, following the same allocation procedure as intercropping assuming that sole soybean is sequentially allocated to the highest yielding sites. We find that 50% soybean self-sufficiency could be reached by cultivating sole soybean on 7.1 Mha, representing 6.0 Mha less compared to intercropping’ requirements (i.e., 13.1 - 7.1 Mha). Based on our maize yield projections, growing sole maize on the 6.0 Mha of land spared under sole soybean scenario could produce 39.0 Mt of maize, i.e., 39% less than the 63.7 Mt produced under intercropping for the same total area of 13.1 Mha (Table 1). To match the total co-production of 18.2 Mt of soybean and 63.7 Mt of maize achieved on 13.1 Mha of maize-soybean intercropping, an additional 3.4 Mha of sole maize would be required, increasing the total land required by sole cropping to 16.5 Mha. These results indicate that intercropping can achieve similar co-production of both crops using 21% less land compared to sole cropping (i.e., [16.5 – 13.1] / 16.5), while covering 50% EU soybean self-sufficiency. Similar results are observed for soybean self-sufficiency levels of 25% and 75%, for which intercropping allowed for 21% and 20% land saving compared to sole cropping, respectively (Table 1).

Land saving allowed by intercropping slightly decreases in simulations considering high N inputs and strong overlap between soybean and maize growing periods, two factors which were found to reduce maize-soybean intercropping efficiency.^22^ Assuming N fertilization rates equivalent to those of maize in 2020 in the EU,^33^ pLERs values varied from 0.45 and 0.59 for soybean and 0.77 and 0.79 for maize across the EU,^22^ respectively (Supplementary Figure 11). In this case, intercropping uses 19% and 21% less land compared to sole cropping to achieve similar co-production of both crops considering the targets of 25% and 50% soybean self-sufficiency in the EU (Supplementary Table 4). Similarly, we obtain land savings of 16% and 18% when targeting 25% and 50% soybean self-sufficiency in a scenario where both species are sown and harvested at the same time (i.e., characterized by a temporal niche differentiation [TND] of 0), corresponding to pLERs values of 0.53 and 0.76 for soybean and maize,^22^ respectively (Supplementary Table 5).

### Potential contribution of a large-scale deployment of intercropping to EU’s self-sufficiency

Our findings suggest that growing maize-soybean intercropping in soybean-suitable area every four years could substantially contribute to both soybean and maize self-sufficiency in the EU, while improving land use efficiency.

In an optimistic scenario where the total intercropping area could be as high as 25 Mha, our estimations indicate that maximum self-sufficiency levels of 91% for soybean and at least 145% for maize could be achieved with a conventionally managed intercropping system (i.e., one-in-four years cultivation, with a soybean pLER of 0.56 and a maize pLER of 0.79) (Figure 3). In these conditions, 33.1 Mt of soybean and 123.6 Mt of maize could be produced, representing a total coproduction of 156.7 Mt (Table 1). Considering the same cropland area of 25 Mha, we estimate that allocating 14.8 Mha to sole soybean cultivation could achieve 100% soybean self-sufficiency in the EU. However, growing maize as sole crop on the remaining 10.2 Mha could result in 56% lower maize production compared to intercropping. Maize-soybean intercropping could thus benefit to EU’s self-sufficiency not only in soybean, but also in maize.

**Figure 3.**
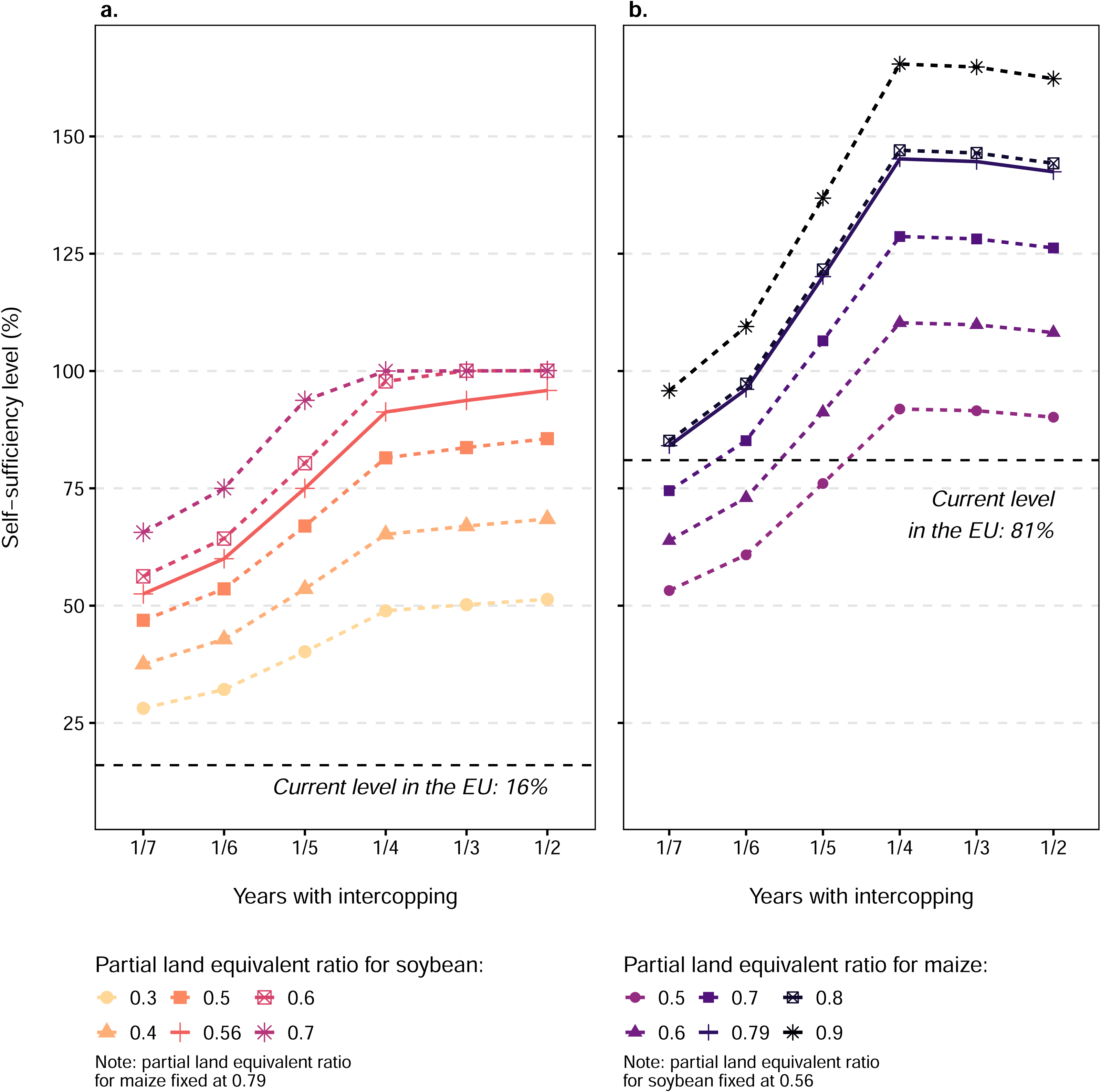
Levels of soybean (a) and maize (b) self-sufficiency in the European Union (EU) achieved from intercropping for different assumptions of partial land equivalent ratio (pLER) and crop return frequency. Crop frequencies of one year in seven, six, five, four, three, or two correspond to allocating maize-soybean intercropping on 14%, 16%, 20%, 25%, 33%, or 50% of cropland area in each grid-cell. Intercropping was first allocated to soybean highest-yielding grid-cells, and in all cases total intercropping area was constrained to not exceed 25% of croplands in the EU (i.e., 25 Mha). Squares represent the results obtained with average productive performances of maize-soybean intercropping systems (i.e., pLER of 0.56 for soybean and 0.79 for maize) estimated in a previous meta-analysis (Xu et al., 2020).

The sensitivity of these results to variations in intercropping’s productive performances, return frequency, and area of cultivation is also explored. Provided that 25 Mha is allocated to intercropping, we estimate that 29-100% soybean and 53-165% maize self-sufficiency are potentially achievable through intercropping, depending on return frequency and assumed pLER values (Figure 3). In particular, more than 53% and 84% self-sufficiency for each crop are achievable when pLER values remain equal to 0.56 for soybean and 0.79 for maize, respectively. Similar levels of self-sufficiency are achieved after incorporating the effects of N inputs and TND on the pLERs. In the analysis which incorporates the effect of N fertilization rates, the potential self-sufficiency rates range from 49 to 90% for soybean and 83 to 143% for maize, respectively, depending on the return frequency. Similarly, 49-90% and 80-138% self-sufficiency for soybean and maize, respectively, could be achieved in a scenario characterized by a TND of zero (i.e., corresponding pLERs values: 0.53 and 0.76), respectively (Supplementary Figure 12). More generally, when maintaining maize pLER at 0.79 while reducing soybean pLER to 0.5, 0.4, or 0.3, a minimum frequency of one year in six, five, or three is required to meet 50% soybean self-sufficiency, respectively (Figure 3a). With a one-in-four years frequency, 48-100% soybean self-sufficiency could be reached when soybean pLER varies from 0.3 to 0.7. It is worth noting that potential self-sufficiency rates achievable with intercropping always exceeds the current level in the EU for soybean, even in the least productive intercropping scenario (i.e., one-in-seven years frequency and 0.3 soybean pLER). To exceed the current maize self-sufficiency level in the EU, growing maize-soybean intercropping every six, five, and four years could be required with maize pLER values of 0.5, 0.6, and 0.7, respectively, assuming a 0.56 soybean pLER (Figure 3b).

Sensitivity analyses investigating the effect of variations in pLERs values for soybean (from 0.3 to 0.7, by 0.01-increment) and maize (from 0.5 to 0.9, by 0.01-increment) show a trade-off between the pLER values and the total allocated surface: as intercropping becomes more efficient for soybean production compared to sole crop (i.e., as soybean pLER increases), less land is required to meet the soybean self-sufficiency target, thus reducing the production of companion maize associated to soybean in intercropping (because the area allocated to intercropping decreases). To illustrate this, Supplementary Figure 13 shows the results of these sensitivity analyses when targeting 25% of soybean self-sufficiency (i.e., 9.1 Mt of soybean) in the EU. This level of soybean self-sufficiency is met as soon as soybean pLER is higher than 0.3 (panel a of Supplementary Figure 13). Consequently, maize production and intercropping area decrease as the value of soybean pLER increases, and this area reduction leads to a decrease of the total maize produced by the intercrops if the maize pLER remains unchanged. For example, when the soybean pLER increases from 0.3 to 0.56, the area required to produce 9.1 Mt of soybean decreases from 12.2 to 6.3 Mha, respectively (panel c of Supplementary Figure 13), reducing the maize production on the intercropping area from 58.9 to 28.8 Mt when assuming a fixed value of maize pLER of 0.79. Increasing the maize pLER mitigates the decrease in the amount of maize produced by intercropping. When fixing the soybean pLER at 0.3, the minimum maize pLER values required to reach 50% and 75% of maize self-sufficiency levels are 0.58 and 0.86, respectively (panel d of Supplementary Figure 13).

Similarly, the level of 50% soybean self-sufficiency is reached as long as soybean pLER is higher than 0.31, assuming that 25 Mha is allocated to maize-soybean intercropping (panels a and c of Supplementary Figure 14). Interestingly, 50% soybean self-sufficiency and at least 100% maize self-sufficiency could be simultaneously achieved with a maize pLER of 0.79 as soon as soybean pLER remains lower than 0.41. When soybean pLER exceeds this threshold, less land is required to reach 50% soybean self-sufficiency and the area allocated to intercropping decreases. In response, maize self-sufficiency does not exceed 100% and could drop to 75% under highest soybean pLER values (panels b-c of Supplementary Figure 14). When targeting 75% or 100% soybean self-sufficiency using maize-soybean intercropping, one limiting factor was the maximum area allocated (Supplementary Figures 15-16). Within the constraint of 25 Mha (i.e., 25% of European cropland) of maize-soybean intercropping, a soybean pLER of 0.46 is required to meet three-quarters of the EU’s demand in soybean, while achieving full soybean self-sufficiency could require a minimum pLER of 0.62.

Our main results are obtained assuming that soybean could be cultivated in all areas with a soybean yield higher than 1 t.ha^-1^. If we restrict soybean cultivation to the 25% most productive sites only (i.e., sites with soybean yield ≥2.6 t.ha^-1^), 11-98% soybean self-sufficiency could be achieved, depending on cropping frequency and soybean pLER (Supplementary Figures 17-18). On the opposite, extending soybean cultivation in several third countries and candidates for accession to the EU (Azerbaijan, Belarus, Norway, Switzerland, United-Kingdom, Albania, Bosnia and Herzegovina, Georgia, Moldova, Montenegro, North Macedonia, Serbia, Türkiye, Ukraine, and Kosovo) improves the potential soybean self-sufficiency levels attainable through intercropping (43-100%), especially in case of low return frequency (i.e., one-in-seven to one-in-four years cultivation) (Supplementary Figure 19). In this case, intercropping area shifts from northern and southern Europe to eastern Europe, especially in Ukraine, compared to the allocations performed for the EU only (Supplementary Figures 20).

## Discussion

This study estimates the benefits of intercropping to improve EU’s self-sufficiency in soybean while maintaining a high production of maize, and quantifies the land saved by this strategy over sole cropping at a continental scale. It extends the results from previous meta-analysis by taking into account spatial variability of maize and soybean productivity in the EU. Our results suggest that growing maize-soybean intercropping in areas most suited for soybean cultivation has the potential to enhance EU soybean and maize self-sufficiency, thus substantially reducing Europe’s dependency to imported supplies. Based on geographical patterns of sole crops’ yields, a one-in-four-year return frequency, and assuming that 1 ha of intercropping produces as much as 0.56 ha of soybean and 0.79 ha of maize in pure stands,^22^ 13.1 Mha of intercropping could cover 50% and 75% of the EU’s consumption in soybean and maize (i.e., 18.2 Mt of soybean and 63.7 Mt of maize), respectively. Additionally, maize-soybean intercropping emerges as a powerful strategy for improving land use efficiency in the EU. Intercropping outperformed sole cropping in maximizing maize-soybean co-production, allowing substantial land savings. For instance, to simultaneously produce 18.2 Mt of soybean and 63.7 Mt of maize, intercropping required 3.4 Mha (21%) less land than sole cropping, corresponding to an area slightly higher than the combined agricultural areas of Belgium and the Netherlands (1.4 and 1.8 Mha, respectively).^32^ This land saving estimate differs from the value obtained by direct extrapolation from the pLER values. Indeed, assuming average soybean and maize pLERs of 0.56 and 0.79,^22^ respectively, expected land savings based on field-scale experiments could be 26% as compared to the 20-21% estimated here. This difference demonstrates that pLER values derived from field-scale experiments cannot be robustly extrapolated to continental-scale land-use efficiency, because geographic variation in actual yields of maize and soybean need to be accounted for (see Supplementary Note 1 for a detailed explanation of how spatial yield variations affect estimates of land savings).

Previous research has shown that the productive performances of intercropping are highly variable, with LER values ranging from less than 0.5 to more than 2.0 for maize-soybean intercropping on the global scale.^22^ While intercropping scenarios characterized by pLERs values higher than 0.5 for soybean and 0.7 for maize lead to higher improvements in self-sufficiency and land savings, our results suggest that intercropping maize and soybean in the EU remains an interesting option to improve self-sufficiency compared to sole crops even with lower pLERs. As an example, the level of 50% soybean self-sufficiency is reached as long as soybean pLER is higher than 0.3, assuming that 25 Mha is allocated to maize-soybean intercropping (Figure 3). In this study, intercropping’s contribution to maize and soybean self-sufficiency slightly decreased with higher N inputs or when maize and soybean were grown during the same period, but still strongly improved self-sufficiency compared to the current EU level. These results highlight the importance of identifying combinations of environmental conditions and management practices that maximize the benefits of this strategy. Many factors affecting the performances of intercropping have been identified in previous research. For example, results from a meta-analysis of field trials conducted in China^34^ suggest that increased sunshine hours and temperatures could reduce the performance of maize-soybean intercropping compared to sole crops. It has been shown that due to interspecific competition, soybean often underperforms relative to maize in intercropping,^35^ particularly under increased nitrogen fertilization^22^ and nutrients availability.^34^ However, wider strip distance,^36^ moderate maize planting density, or shorter growth periods’ overlap^34^ could contribute in reducing the competition between species in intercrops. In Europe, the performance of cereal-legume mixtures is also affected by climate conditions, especially rainfall, and farming practices such as tillage, and nature and quantity of applied fertilizers.^37^ Given the limited information on the interactions between soybean productivity in intercropping and other farming practices in Europe, additional trials are required to refine local estimations of maize-soybean intercropping performances and to help identifying the best combinations of field configuration, fertilization, and water management which maximizes the productivity of intercropping at the farm level.

Although several studies suggest that deploying intercropping in Europe could provide agroecological benefits while maintaining crop productivity,^38^ its development is hampered by numerous barriers^39,40^ throughout the food value chain, but predominantly faced by farmers.^41^ Tackling the farmer-specific barriers is necessary but insufficient to promote widespread adoption across the EU.^39,41^ To comply with grain collectors and processors purity standards, intercropping requires specific machinery for sowing, harvesting, and sorting of grains of the different crops. Machinery is already available for intercropping, especially in China,^42^ but further development is required to facilitate their large-scale deployment. In addition, adapted irrigation systems and crop protection practices still need to be identified, also according to intercropping design. Large-scale introduction of intercropping could create a substantial market, driving the development of affordable, specialized machinery. However, such equipment will not be sufficient to make intercropping a viable system if purity standards are not adapted to intercropping products. Given the systemic nature of these barriers, involving all stakeholders across the value chain is essential for designing effective policies to unlock the full potential of legume-cereal intercropping in Europe.^40^ On the positive side, since genetically modified crops are only marginally grown in the EU, the soybean produced in EU will be non-genetically modified, and will therefore address a specific market, different from that for genetically modified soybean from North and South America. Additionally, adopting maize–soybean intercropping could alter crop sequences, likely reducing the temporal frequency of some crops (e.g., maize) while increasing that of others (e.g., soybean). Such increases in temporal crop diversity within rotations offer multiple agronomic benefits, including improved control of pests, diseases, and weeds, as well as enhanced nitrogen management (particularly when including legume crops) with positive effects on crop yields.^43,44^ However, quantitative evidence on the effects of crop sequences that include maize–soybean intercropping remains limited. Although a few studies exist,^45^ the available data are currently insufficient to support robust assessments.

One strength of this study is the use of a machine learning model to model soybean and maize productivity trained on a global historical yield dataset.^46^ Thanks to their greater flexibility, machine learning algorithms often outperform traditional methods (e.g., linear regression or process-based models) to forecast yields of crops from climate predictors, especially soybean^47,48^ and maize.^49^ For soybean, a limitation of the model is the low number of observations collected in Europe as few data points were available in Europe to train the model for soybean, which may affect prediction accuracy. Our results highlight that maize-soybean intercropping performs well under current climate, but the suitability of this cropping strategy in future climate conditions has not been examined in our analysis. Previous studies^12,14^ suggest that by 2050, climate change may shift soybean-suitable area in Europe, with northern regions becoming more favorable and lower latitudes increasingly less suitable.^50,51^ Overall, it has been shown soybean production in Europe is likely to increase under various climate change scenarios compared to historical conditions.^12,51^ On the contrary, maize yields are expected to decline, with an estimated 10% loss per degree Celsius increase if no adaptation strategies are implemented.^49^ However, maize was found to be highly responsive to adaption strategies, such as irrigation and cultivar adaptation, which could mitigate future yield losses, especially in higher latitudes like Europe.^49,52^ More research is needed to assess the robustness of our findings in light of these potential changes in the spatial yield patterns of soybean and maize. With the possible reduction of the lengths of growing periods in the context of climate warming, the cultivation of several crops during one single growing season (i.e., relay or double cropping) might become more plausible in some regions.^53,54^ However, a comprehensive analysis of the relative interest of these practices compared to intercropping is lacking. In absence of experimental datasets that explicitly explore variations in intercropping efficiency as a function of weather conditions, it was not possible to predict the effect of environmental factors on pLERs and, thus, take the potential spatial variability of pLERs into account. The establishment of a shared, standardized, and open-access global database on intercropping systems—including detailed information on management, environmental conditions, and LER outcomes—could represent a major advance to address this issue.

Assuming that 1 ha of intercropping produces as much as 0.56 ha of sole soybean and 0.79 ha of sole maize, our results suggest that 13.1 Mha of intercropping are required to meet 50% soybean self-sufficiency in the EU (Table 1). This area is only 30% larger than the 10 Mha currently devoted to the cultivation of maize and soybeans in the EU (Supplementary Table 2). However, achieving higher soybean self-sufficiency levels through intercropping expansion could have a more important impact on land use in the EU. Our findings suggest that, in the same conditions of land use efficiency (pLER of 0.56 and 0.79 for soybean and maize, respectively), 25 Mha of maize-soybean intercropping could be required to achieve 91% soybean self-sufficiency. This could be achieved by growing maize-soybean intercrops on the current harvested area of maize and soybean in the EU plus an additional 15 Mha currently cultivated by other crops, mostly wheat, barley rapeseed and sunflower (Supplementary Table 3). In this scenario, the pressure on cropland will increase, but it is difficult to determine which crops are most likely to be replaced by corn-soybean intercrops due to the influence of numerous factors, including future supply-demand balances and future commodity prices.^55^

Our results suggest that the widespread adoption of maize-soybean intercropping in the EU could improve soybean self-sufficiency (25-91%, depending on the considered scenario) but also contribute to cover maize demand in the EU (50-145%, depending on the considered scenario). For maize, the levels attained could even be higher considering the maize produced outside of the intercropping area, thus increasing EU’s maize self-sufficiency and exports. However, our sensitivity analyses suggest that the design of intercropping systems could be adjusted (especially by tuning intercropping return frequency, relative crop densities, and maize pLER) to limit maize overproduction while still achieving high levels of soybean self-sufficiency. Although maize-soybean intercropping offers advantages in terms of productivity,^56^ a large-scale deployment of this type of intercrop could compete with other crops that play an important role in feed and food production, such as barley and sunflower. To limit this problem, it could be advisable to diversify the types of intercropping and consider the use of double cropping systems, i.e., systems with two crop harvests per year, in regions where this system could be implemented.^12,48,57^ Potential production from other intercropping systems and double cropping systems deserves further research.

An alternative strategy to reduce EU dependency on soybean imports is to decrease the soybean demand. This could be achieved by growing other legume crop species (e.g., broad bean, pea, and lupin) as feeds^16^ or by switching to a less meat-based diet with lower meat consumption by humans.^5^ The latter was found to reduce the demand for soybean-based protein in the EU through lower meat consumption, and free up land to produce more legume crops for human consumption rather than livestock feed. In the future, an integrated strategy combining both an increase in supply through intercropping and a reduction in demand through a partial change in diet could enable Europe to reduce its dependence on imports of agricultural commodities without too much impact on its cultivated areas.

## Methods

### Yield projections in the EU

We developed models to forecast soybean and maize grain yields (at standard moisture percentage) in the EU – referring to the EU from February 1^srt^ 2020, when the United Kingdom withdrew – as a function of climate conditions and water management. For both crops, we built a training dataset including gridded data on historical worldwide yields collected in major producing areas,^46^ irrigated fraction from the SPAM2010 database,^58^ daily minimum and maximum temperature (°C), total precipitations (mm), solar radiations (MJ.m^-^²), evapotranspiration (mm.day^-1^), and vapor pressure deficit (kPa) from the ERA5-Land database.^59^ For each site, we expressed annual yield data relatively to the expected value at the most recent year available (i.e., 2016). This detrending procedure allowed to remove the systematic temporal component of yield that is attributed to technological progress (e.g., genetic improvement, management, inputs) and to focus on the interannual variability around that trend, which is typically interpreted as being driven by climate, weather, soil, or other non-progress-related factors. Further details on data pre-processing steps are provided in Supplementary Note 3. The total number of sites (resp. number of sites-year) included in the soybean and maize datasets reached 3,286 (116,516) and 2,064 (72,964), respectively. In production sites, detrended yields ranged from 0.1 to 6.7 t.ha^−1^ for soybean and from 0.5 to 20.0 t.ha^-1^ for maize (Supplementary Figure 21). The minimum yields observed in the train dataset for major regions producing soybean with climatic conditions comparable to those of the EU (i.e., Italy and the US) and major maize-producing regions in the EU were 1.0 and 1.3 t.ha^-1^, respectively. Mean (standard deviation) yield, representing the average (standard deviation) of values rescaled to the technological baseline of 2016 and independent of the original calendar years of observation, was 2.6 t.ha^−1^ (1.1 t.ha^−1^) for soybean and 9.5 t.ha^−1^ (3.8 t.ha^-1^) for maize. Exclusively rainfed soybean or maize were grown in 45% and 24% of sites, respectively. Within sites with fractional area of irrigated soybean or maize exceeding 0%, irrigated soybean and maize represented 1-22% and 1-30% of cultivated area of each crop, respectively.

We tested different approaches to predict soybean and maize yields (Supplementary Note 4). The choice of the models was guided by previous work comparing various techniques to aggregate climate predictors and various algorithms to predict soybean yield.^12,47^ These studies showed that the first two principal components (PCs) were able to capture large variations in monthly climate averages and using these PCs in a random forest algorithm provided highly accurate predictions of soybean yields in rainfed conditions. Following this methodology, we built distinct predictive models for both crops, as well as variants based on different climate data aggregation methods (i.e., monthly average, seasonal average, or a higher number of PCs) for comparison. For each crop, the model based on the first two PCs showed the best transferability in time and space according to two cross-validation procedures (Supplementary Note 4). Based on this approach, we projected the annual yields of rainfed soybean and maize in pure stands in 2699 sites EU over 2000-2023 (i.e., 64776 site-year) using climate data from the ERA5-Land database^59^ as predictors. Except for soybean yields ≥3.0 t.ha^-1^ and maize yields ≥7.0 t.ha^-1^, projections were not sensitive to the models selected (Supplementary Figure 22).

### Performance of intercropping to maximize soybean and maize co-production in the EU

We estimated the yield of both crops when cultivated in intercropping by multiplying the projections of yields in pure stands and rainfed conditions by their respective values of pLER (i.e., 0.56 and 0.79, respectively) obtained from a meta-analysis of soybean-maize intercropping trials.^22^ Results of this meta-analysis were estimated using 654 field observations collected in 108 sites worldwide, with environmental conditions partially representative to those in the EU (Supplementary Figures 23-24). Although this meta-analysis also provides an estimation based on data collected in Europe specifically, we choose to use the global estimate, which is slightly less optimistic (i.e., lower pLER), preventing us to overestimate the productivity of intercropping. We then estimate the production of soybean and maize as the product of predicted intercropping yields, site-specific cropland area (corresponding to “arable lands and permanent crops” based on FAO’s definition) from the SASAM dataset,^60^ and a coefficient representing the return frequency of intercropping in crop sequences. We set this coefficient at 1/4, corresponding to a one-in-four years return frequency to limit the risk of pests and diseases^30,31^ and which is equivalent to allocating each year 25% of the cropland area in each site.

Based on these estimates, we examined the potential of intercropping to improve EU’s soybean self-sufficiency while limiting competition with other crops by performing allocation scenarios of maize-soybean intercropping across the EU. Our allocation process was guided by three assumptions: (1) maize-soybean intercropping is preferentially implemented in the cropland areas with the highest projected soybean production but (2) excluding sites with unsuitable conditions for soybean production (i.e., projected yields below 1 t.ha^-1^, corresponding to the minimum observed in the training dataset from the US and Italy; Supplementary Figure 21); (3) we assumed that intercropping could first replace existing maize and soybean areas (i.e., 10 Mha including 1.0 Mha of soybean and 8.9 Mha of maize, 2018–2022 averages).^6^ If the soybean self-sufficiency targets were not met, additional cropland could be allocated up to 15 Mha to limit competition with other crops. In that case, the total intercropped area at the scale of the EU is capped at 25 Mha, an area equivalent to 23% of the EU’s cropland (i.e., 109 Mha)^26^ and comparable to the current European wheat area in 2020.^61^ In the area allocated to maize-soybean intercropping, the minimum maize yields projections (ranging from 3.7 to 4.0 t.ha^-1^ depending on the considered scenario) were three-to-four times higher than the minimum yield of 1.3 t.ha^-1^ observed in our train dataset of maize-producing regions in the EU (Supplementary Figure 21). Considering this, no additional criterion was considered for maize suitability in the allocation procedure.

For each allocation scenario, maize-soybean intercropping was sequentially allocated to highest yielding sites, ranked by soybean productivity (i.e., projected soybean yield multiplied by the cropland area in each site), until 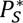 was reached or until the total area allocated to intercropping reached 25 Mha. This allocation yielded to simultaneously produce 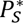 Mt of soybean and 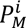 Mt of maize, co-produced through intercropping. The total co-production (i.e, 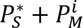) was calculated and compared to the one obtained with a sole cropping strategy applied over the same total cropping area (i.e., the area required to reach 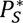 with intercropping). For the sole cropping strategy, we first allocated sole soybean to the highest yielding sites until reaching 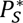 or 25 Mha. Second, we allocated sole maize to the remaining sites and computed the resulting maize production in sole crop maize 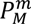. Note that, in this approach, the sum of the sole soybean area and the sole maize area is equal to the intercropping area obtained with the intercropping allocation strategy previously described.

Since both allocation procedures were designed to produce 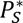 Mt soybean, the most efficient strategy among intercropping (i.e., simultaneous cultivation of maize and soybean in the same area) or sole cropping (i.e., separate cultivation in different areas) was the one leading to the largest production of maize in the region of interest. We performed four distinct allocation scenarios considering values of 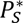 of 9.1, 18.2, 27.2, and 36.3 Mt, equivalent to soybean self-sufficiency levels of 25%, 50%, 75%, or 100% in the EU, respectively, based on the average EU soybean consumption of 36.3 Mt over the 2018–2022 period.^6^ For each scenario, we estimated the areas required by intercropping and sole cropping. Based on these estimates, land saving allowed by the most efficient strategy over the least efficient one was estimated as the difference between the area required by the least efficient strategy to reach the same level of co-production than the most efficient strategy and the area required by the most efficient strategy, divided by the area required by the most efficient strategy. Further details regarding the procedure to allocate and compare sole and intercropping strategies are provided in Supplementary Note 2.

### Sensitivity analyses

To assess the sensitivity of our results to cropping frequency, the main analysis (i.e., pLER values of 0.56 for soybean and 0.79 for maize, respectively) was repeated in scenarios specified by combining different values of crop frequencies (one-in-seven to one-in-two years, by yearly increment), soybean pLER (0.3 to 0.7, by 0.1-increment) and maize pLER (0.5 to 0.9, by 0.1-increment). These values of pLERs corresponded to LER ranging from 0.8 to 1.6, in the range of the possible values of maize-soybean LER estimated by Xu et al.^22^

The results of Xu et al.^22^ show that soybean pLER vary according to fertilization practices, especially N inputs but also depending on the overlap between growing periods of maize and soybean, characterized by the TND. High TND values (i.e., close to 1) correspond to small overlap, where the second species is sown just before the harvest of the first crop (e.g., relay intercropping), whereas a TND of zero represents systems in which both species are sown and harvested simultaneously. To evaluate the effects of N inputs and TND on values of partial land equivalent ratios (pLERs) of soybean and maize, we used the regression models published by Xu et al.^22^ to compute less optimistic pLER values (i.e. lower) assuming high N fertilizer doses and low TND and assessed the robustness of our conclusions considering these lower pLER values. First, to take into account the spatial variability in N management in the EU, we computed site-specific soybean and maize pLERs according to local N fertilizer application rates of 2020 provided by the NPKGRIDS gridded dataset^33^ using published regression model.^22^ To compute less optimistic pLER values (i.e. lower) assuming high N fertilizer doses, we considered that the applied N fertilizer rates were those of maize, which receive higher amounts of N. Supplementary Figure 10 shows the applied rates, as well as derived pLER values. Second, we assumed that both species had identical growing periods (i.e., TND=0), which led to pLER values of 0.53 and 0.76, respectively. Further details regarding these sensitivity analyses are provided in Supplementary Note 5.

Additionally, for the different soybean self-sufficiency levels considered in this study (i.e., 25%, 50%, 75%, and 100%), we considered scenarios with varying pLER values of soybean (0.3 to 0.7, by 0.01-increment) and maize (0.5 to 0.9, by 0.01-increment) to increase the range of possible performances of intercropping over sole cropping. We estimated the minimum pLERs conditions simultaneously satisfying each soybean self-sufficiency levels, as well as 25%, 50%, 75%, and 100% maize self-sufficiency, given a one-in-four-year return frequency and limiting the area allocated to intercropping to 25% of the cropland area in the EU.

To explore the impact of concentrating intercropping in the high-yield sites, allocation procedures were repeated for sites where the average soybean yield exceeded 2.6 t.ha^-1^ (instead of 1 t.ha^-1^) representing the 75^th^ percentile of projected soybean yields. The impact of expanding soybean cultivation in several third countries and candidates for accession to the EU was explored as well.

## Data Availability Statement

The processed data are available in the “Maize-Soybean Intercropping in Europe” repository, available at: https://github.com/MathildeChen/Maize_Soybean_Intercropping_Europe (DOI: 10.5281/zenodo.20794811).

## Code Availability Statement

All relevant R (version 4.3.1 http://www.r-project.org) scripts and documentation are available via the project repository on Github at https://github.com/MathildeChen/Maize_Soybean_Intercropping_Europe.

## Funding Statement

The work of MC, NG, and DM was supported by the CLAND convergence institute (16-CONV-0003) funded by the French National Research Agency (ANR).

## Author Contributions Statement

Concept and design: MC, NG, DM; Acquisition of data and statistical analysis, including the production of results, figures, and tables: MC; Interpretation: MC, NG, DM; Drafting of the manuscript: MC; Critical revision of the manuscript: MC, NG, DM; Obtained funding: NG, DM; Administrative, technical, or material support: DM; Supervision: NG, DM.

All authors approved the submitted manuscript and all revised versions. Each author accepts personal responsibility for their contributions and agrees to ensure that any questions concerning the accuracy or integrity of the work are appropriately investigated, resolved, and that the outcomes of such investigations are properly documented in the scientific literature.

## Competing Interests Statement

The authors declare no competing interests.

## Notes

https://github.com/MathildeChen/Maize_Soybean_Intercropping_Europe/tree/main

### Competing Interest Statement

The authors have declared no competing interest.

### Summary of Updates

Revisions adressing editorial requests. The number of words in the Abstract was reduced. Figure 3 has been revised so that symbols match between the legend and the figure. All Supplementary figures and tables are now numbered and labeled. Maps in Supplementary Figures 1, 10, 11, 18, 20, 22, 23a, and 24a now show longitudes and latitudes. Supplementary Figure 22 has been modified using the colorblind friendly color palette.The references are now fully indicated.

