## Supplementary Information for "High potential contribution of intercropping to soybean and maize self-sufficiency in Europe"

* Corresponding authors:

**List of Supplementary Notes**

Supplementary Note 1. Productive performance of intercropping in case of spatial variability of yield ... 5

Supplementary Note 2. Procedure for soybean and maize allocation in Europe ... 8

Supplementary Note 3. Training dataset constitution and pre-processing steps of data analyses 10

Supplementary Note 4. Assessment of the predictive performance of several crop yield forecasting models ... 14

Supplementary Note 5. Computation of partial land equivalent ratios of soybean and maize for sensitivity analyses including the effect of fertilization and temporal niche differentiation on partial land equivalent ratios ... 15

**List of Supplementary Figures**

Supplementary Figure 1. Geographical distribution of sites included in soybean (a) and maize (b) the training dataset. Soybean and maize training datasets included 116,516 and 72,964 unique combinations of sites and years, respectively. Base map based on Natural Earth data, created using the R package rnaturalearth. ... 16

Supplementary Figure 2. Scatterplots of soybean (a) and maize (b) yields observations compared to the best model predictions estimated by cross-validation on sites (top panel) or on years (bottom panel). ... 17

Supplementary Figure 3. Distribution of the residuals (predictions - observation) of the best models forecasting soybean (a) and maize (b) yields, estimated by cross-validation on sites (top panel) or on years (bottom panel). ... 18

Supplementary Figure 4. Importance of predictors in the random forest models showing best performance to predict soybean (a) and maize (b) yields in the training datasets. ... 19

Supplementary Figure 5. Contribution of climate data to the first principal component derived from principal component analysis applied on monthly averages (a) and partial dependency plot associated with each predictor (b) in the best model predicting *soybean* yield. ... 20

Supplementary Figure 6. Contribution of climate data to the second principal component derived from principal component analysis applied on monthly averages (a) and partial dependency plot associated with each predictor (b) in the best model predicting *soybean* yield. ... 21

Supplementary Figure 7. Contribution of climate data to the first principal component derived from principal component analysis applied on monthly averages (a) and partial dependency plot associated with each predictor (b) in the best model predicting *maize* yield. ... 22

Supplementary Figure 8. Contribution of climate data to the second principal component derived from principal component analysis applied on monthly averages (a) and partial dependency plot associated with each predictor (b) in the best model predicting *maize* yield. ... 23

Supplementary Figure 9. Partial dependency plot associated with irrigated fraction in the best model predicting soybean (a) and maize (b) yield. ... 24

Supplementary Figure 10. Maize-soybean intercropping and sole cropping allocations for 25, 50, 75, and 100% soybean self-sufficiency (i.e., 9.1, 18.2, 27.2, and 36.3 Mt, respectively) in the European Union. ... 25

Supplementary Figure 11. The performances of maize-soybean intercropping according to (a) nitrogen fertilization rates for maize of 2020, represented by the partial land equivalent ratios of (b) soybean and (c) maize. Gridded fertilization rates (0.5°-resolution) were obtained from the

NPKGRIDS dataset (11). Soybean and maize pLERs values were computed based on the equations of Xu et al. (12). Base map based on Natural Earth data, created using the R package rnaturalearth. 26

Supplementary Figure 12. Levels of soybean (a) and maize (b) self-sufficiency in the European Union (EU) achieved from intercropping for different assumptions of partial land equivalent ratios (pLERs) and crop return frequency. 27

Supplementary Figure 13. Characteristics of maize-soybean intercropping allocations in terms of soybean produced (a), maize produced (b), and surface allocated (c) for covering 25% soybean self-sufficiency in the European Union depending on intercropping efficiency (pLER). The minimum efficiency conditions simultaneously satisfying 25% of soybean and 25, 50, 75, or 100% maize self-sufficiency are also shown (d). 28

Supplementary Figure 14. Characteristics of maize-soybean intercropping allocations in terms of soybean produced (a), maize produced (b), and surface allocated (c) for covering 50% soybean self-sufficiency in the European Union depending on intercropping efficiency (pLER). The minimum efficiency conditions simultaneously satisfying 50% of soybean and 25, 50, 75, or 100% maize self-sufficiency are also shown (d). 29

Supplementary Figure 15. Characteristics of maize-soybean intercropping allocations in terms of soybean produced (a), maize produced (b), and surface allocated (c) for covering 75% soybean self-sufficiency in the European Union depending on intercropping efficiency (pLER). The minimum efficiency conditions simultaneously satisfying 75% of soybean and 25, 50, 75, or 100% maize self-sufficiency are also shown (d). 30

Supplementary Figure 16. Characteristics of maize-soybean intercropping allocations in terms of soybean produced (a), maize produced (b), and surface allocated (c) for covering 100% soybean self-sufficiency in the European Union depending on intercropping efficiency (pLER). The minimum efficiency conditions simultaneously satisfying 100% of soybean and 25, 50, 75, or 100% maize self-sufficiency are also shown (d). 31

Supplementary Figure 17. Level of soybean self-sufficiency in the European Union (EU) achieved by intercropping on areas showing soybean productivity equal or higher than 1 t.ha$^{-1}$ (left panel) or 2.6 t.ha$^{-1}$ (right panel), in several scenarios of partial land equivalent ratio (pLER) and crop return frequency. 32

Supplementary Figure 18. Geographical allocation of maize-soybean intercropping in areas showing soybean productivity equal or higher than 1 t.ha$^{-1}$ or 2.6 t.ha$^{-1}$ (a) and soybean yields projected in corresponding sites (2000-2023) (b). 33

Supplementary Figure 19. Level of soybean self-sufficiency in the European Union (EU) achieved from intercropping on areas showing soybean productivity equal or higher than 1 t.ha$^{-1}$ in the EU exclusively (left panel) or extended to neighboring countries (right panel), in several scenarios of partial land equivalent ratio (pLER) and crop return frequency. 34

Supplementary Figure 20. Geographical allocation of maize-soybean intercropping in the European Union exclusively (EU) or extended to neighboring countries (EU extended) (a) and soybean yields projected in corresponding sites (2000-2023) (b). 35

Supplementary Figure 21. Soybean (a) and maize (b) yields distribution from 1981 to 2016 in sites located in producing areas included in the training dataset. 36

Supplementary Figure 22. Projected yields of soybean and maize in the European Union according to four different predictive models. 37

Supplementary Figure 23. Location (a), average temperature and total precipitations during soybean growing period over 2000-2023 (b) in the experimental sites used in the meta-analysis of Xu et al. (12) and in the sites where soybean yields are projected in the present study. 38

Supplementary Figure 24. Location (a) and distribution (b) of Köppen-Geiger zones among the sites included in the meta-analysis of Xu et al. (12) [MA] and in the sites where soybean yields are projected in the present study [EU]. 39

Supplementary Figure 25. Variance explained by the principal components derived from principal component analysis applied on each climate variable in the soybean (a) and maize (b) training dataset. 40

**List of Supplementary Tables**

Supplementary Table 1. Nash-Sutcliffe efficiency of climate-based models predicting soybean and maize yields estimated by cross-validation on years, sites, and on average. 41

Supplementary Table 2. Soybean and maize food balance, self-sufficiency rate, areas, and productivity in the European Union. 42

Supplementary Table 3. Area of major crops in regions with high suitability of soybean cultivation in the European Union. 43

Supplementary Table 4. Soybean and maize coproduction and surface requirement for 25, 50, 75, and 100% soybean self-sufficiency in the European Union (EU) achieved by intercropping or sole cropping *according to local nitrogen fertilization rates in the EU*. 44

Supplementary Table 5. Soybean and maize coproduction and surface requirement for 25, 50, 75, and 100% soybean self-sufficiency in the European Union (EU) achieved by intercropping or sole cropping *assuming that maize and soybean are grown during the same period* 45

### Supplementary Note 1. Productive performance of intercropping in case of spatial variability of yield

As some areas are more productive than others, the allocation of the crop species in the most productive areas may make sole cropping systems more productive than expected according to the land equivalent ration (LER). Although the spatial variability might be relatively low at the field scale, this is an important aspect to consider at large scales.

In the present study, we consider a decision problem combining two distinct objectives, namely:

- The first objective is to produce a target quantity $P_1^*$ of a given crop species (crop 1) in a given region (e.g., country, continent). For example, $P_1^*$ could represent a quantity of crop 1 product corresponding to a high level of self-sufficiency in a given country.
- The second objective is to produce as much as possible of a second crop species (crop 2) in the same region in addition to crop 1. If $P_2$ is the quantity of crop 2 produced, the second objective is thus maximizing $P_2$ considering that a quantity $P_1^*$ should be produced in the region of interest.

In order to achieve these objectives, we compare the performance of two production strategies in order to choose the best one, namely

(i) intercropping of crop 1 and crop 2 (simultaneous cultivation of the two species in the same area),
(ii) crop 1 and of crop 2 cultivated as sole crops (separate cultivation of crop 1 and crop 2 in different areas).

This implies that the best strategy is the one leading to the highest value of $P_2$ in the region of interest while ensuring that a quantity $P_1^*$ of crop 1 is produced in the same region.

Below, we show that the standard criterion LER provides the solution of this decision problem only when the yields of crop 1 and crop 2 are homogeneous over space. Otherwise, LER is not sufficient to choose between the two strategies, even if LER is constant over the intercropping area. Another criterion - based on a modified expression of LER - is defined and, based on this new criterion, I show that a LER>1 is not sufficient to ensure that strategy (i) is the best.

#### Production with intercropping

Let note $A^{(I)}$ the area necessary to obtain a quantity $P_1^*$ of crop 1 with the intercropping strategy. This area is expressed as:

$$A^{(I)} = \frac{P_1^*}{Y_1 \ pLER_1} \quad (1)$$

where $Y_1$ is the average yield of crop 1 grown as sole crop on area $A^{(I)}$, and $pLER_1$ is the partial land equivalent ratio (pLER) equal to the average yield of crop 1 in intercropping to the average yield of crop 1 as sole crop.

As crop 2 is grown together with crop 1 in an intercropping system, the production of crop 2 is expressed as:

$$P_2^{(I)} = Y_2 PLER_2 A^{(I)} \quad (2)$$

where $Y_2$ is the average yield of crop 2 grown as a sole crop on area $A^{(I)}$, and $pLER_2$ is the pLER equal to the average yield of crop 2 in intercropping to the average yield of crop 2 as sole crop.

### Production with separate cultivation

Let note $A^{(M)}$ the area necessary to obtain a quantity $P_1^*$ of crop 1 with the second strategy. This area is considered to represent a subarea of $A^{(I)}$, and is expressed as:

$$A^{(M)} = \frac{P_1^*}{\theta_1 Y_1} \qquad (3)$$

where $\theta_1 Y_1$ is the average yield of crop 1 grown as sole crop on area $A^{(M)}$. The coefficient factor $\theta_1$ specifies the difference in average yield of crop 1 between area $A^{(M)}$ and area $A^{(I)}$. If the yield is homogeneous, $\theta_1 = 1$. If yield of crop 1 is higher in $A^{(M)}$, $\theta_1 > 1$. As usually $pLER_1 < 1$, $A^{(M)} < A^{(I)}$. If we assume that the yield of crop 1 is variable over space and that we allocate crop 1 to the most productive lands first, yield of crop 1 is higher in $A^{(M)}$, and thus $\theta_1 > 1$.

In this strategy, crop 2 is not grown together with crop 1, but separately. In order to consider the same total area as the one obtained with the intercropping strategy, the area allocated to crop 2 is set equal to $A^{(I)} - A^{(M)}$. Then, the production of crop 2 is expressed as:

$$P_2^{(M)} = \theta_2 Y_2 \left[A^{(I)} - A^{(M)}\right] \qquad (4)$$

where $\theta_2 Y_2$ is the average yield of crop 2 grown as sole crops on area $A^{(I)} - A^{(M)}$. The coefficient factor $\theta_2$ specifies the difference in average yield of crop 2 between area $A^{(I)} - A^{(M)}$ and area $A^{(I)}$. Again, if the yield is homogeneous, $\theta_2 = 1$. When the yield of crop 2 is higher in $A^{(I)} - A^{(M)}$ than in $A^{(I)}$, $\theta_2 > 1$ and $\theta_2 < 1$ otherwise.

### Comparison of the two strategies

By definition, both strategies lead to the same level of production of crop 1, i.e., $P_1^*$. However, the production of crop 2 is different. According to eqs.(2) and (4), the ratio between $P_2^{(M)}$ and $P_2^{(I)}$ is expressed as:

$$R = \frac{P_2^{(M)}}{P_2^{(I)}} = \frac{\theta_2 Y_2 [A^{(I)} - A^{(M)}]}{Y_2 pLER_2 A^{(I)}} = \frac{\theta_2 [A^{(I)} - A^{(M)}]}{pLER_2 A^{(I)}} = \frac{\theta_2}{pLER_2}\left[1 - \frac{A^{(M)}}{A^{(I)}}\right]$$

According to eqs.(1) and (3), we have:

$$\frac{A^{(M)}}{A^{(I)}} = \frac{\frac{P_1^*}{\theta_1 Y_1}}{\frac{P_1^*}{Y_1\ pLER_1}} = \frac{pLER_1}{\theta_1}$$

Finally, the production ratio *R* for crop 2 is expressed as:

$$R = \frac{\theta_2}{pLER_2}\left[1 - \frac{pLER_1}{\theta_1}\right] \qquad (5)$$

If the average yields in $A^{(M)}$ and $A^{(I)}$ are equal, the expression of the ratio is reduced to:

$$R = \frac{1 - pLER_1}{pLER_2} \qquad (6)$$

In this case, the production of crop 2 is higher in intercropping if the value of *R* given by eq(6) is lower than 1. This is equivalent to $pLER_1 + pLER_2 > 1$, i.e., *LER* >1.

However, if the average yields in $A^{(M)}$ and $A^{(I)}$ are not equal, then $\theta_1 \neq 1$ and $\theta_2 \neq 1$, and the above condition on LER does not hold anymore. In this case, $R < 1$ is equivalent to

$$\frac{\theta_2}{pLER_2}\left[1 - \frac{pLER_1}{\theta_1}\right] < 1$$

and, finally, to:

$$\frac{pLER_1}{\theta_1} + \frac{pLER_2}{\theta_2} > 1 \qquad (7)$$

The condition (7) is satisfied only when $pLER_1$ ($pLER_2$) is large enough compared to $\theta_1$ ($\theta_2$). In this case, $P_2^{(I)} > P_2^{(M)}$. Otherwise, the condition (7) is not satisfied and $P_2^{(I)} < P_2^{(M)}$

**Example**

We consider an area where the average yields of crop 1 (soybean) and crop 2 (corn) as sole crops are equal to $Y_1$=4 and $Y_2$=8, respectively. We also assume that the pLER of the two crops are equal to $pLER_1$=0.5 and $pLER_2$=0.8, respectively (i.e., the average yields of crop 1 and 2 in intercropping are equal to 4*0.5 and 8*0.8, respectively). The target production level for crop 1 in this area is set equal to $P_1^*$=400.

Based on these assumptions, the requested area with the intercrop system is equal to: $A^{(I)} = \frac{400}{4*0.5} = 200$ and the associated production of crop 2 in the intercrop is equal to $P_2^{(I)}$ = 8 * 0.8 * 200 = 1280.

We now assume that the yield of crop 2 is homogeneous in this total area of 200 ha but that the yield of crop 1 is heterogeneous in the same area. Specifically, in the most productive part of the whole area, we assume that the average yield of crop 1 is 50% higher than the average yield of the whole area. This implies $\theta_1 = 1.5$.The requested area to reach the target production level in the monocrop is thus equal to $A^{(M)} = \frac{400}{1.5*4}$ = 66.67. The remaining area available for crop 2 in monocrop is then equal to $A^{(I)} - A^{(M)}$ =200 - 66.67 and the production of crop 2 as sole cropping is equal to $P_2^{(M)} = 8 * (200 - 66.67) = 1066.64$ and is thus lower than $P_2^{(I)}$. In this case, the best option is the intercropping system and we can easily check that the condition (7) is satisfied:

$$\frac{pLER_1}{\theta_1} + \frac{pLER_2}{\theta_2} = \frac{0.5}{1.5} + \frac{0.8}{1} = 1.13.$$

The conclusion is different if the pLER of crop 2 is decreased to 0.6. In this case, $P_2^{(I)}$=8*0.6*200=960, while $P_2^{(M)}$ is unchanged. The best option is thus the system with sole crops with this new setting, and we can easily check that the condition (7) is not satisfied anymore: $\frac{pLER_1}{\theta_1} + \frac{pLER_2}{\theta_2} = \frac{0.5}{1.5} + \frac{0.6}{1} = 0.933$. Nonetheless, LER is still higher than 1 (0.5+0.6=1.1).

**Supplementary Note 2. Procedure for soybean and maize allocation in Europe**

Allocation of soybean ($S$) and maize ($M$) was performed by optimizing two objectives:

1) produce a target quantity $P_S^*$ of soybean in the EU;

2) produce as much as possible of maize in the same area, in addition to soybean.

If $P_{\mathrm{M}}$ is the quantity of maize produced, the second objective is thus maximizing $P_{\mathrm{M}}$ considering that a given quantity $P_S^*$ should be produced in the region of interest.

The performance in achieving both objectives of maize-soybean intercropping ($i$) was compared to those of separate cultivation of both crops in different areas ($m$). The most efficient strategy was the one leading to the highest value of $P_{\mathrm{M}}$ in the region of interest, while ensuring that a quantity $P_S^*$ of soybean is produced in the same region.

Using the projections of yields in the EU, mean productivity of intercropped soybean in the grid-cell $j$, $P_{Sj}^i$, was computed as follow:

$$P_{Sj}^i = \bar{Y}_{Sj} * pLER_S * F * A_j$$

with $\bar{Y}_{Sj}$ the average of soybean yield projections in the grid-cell $j$ over 2000-2023, $pLER_S$ the pLER of soybean, $F$ the crop return frequency, and $A_j$ the available cropland area in the grid-cell $j$. For each grid-cell, cropland area was retrieved from the SASAM dataset (see previous section). The total production of the $N$ most productive pixels for soybean was thus computed as:

$$P_S^i = \sum_{j=1}^{N} P_{Sj}^i = P_S^*$$

where $N$ defined so that $P_1^I$ reaches the considered target production $P_S^*$.

Similarly, the mean productivity of soybean grown in monocrop in each grid-cell $j$ and total productivity of the $Q$ most productive pixels was:

$$P_{Sj}^m = \bar{Y}_{Sj} * F * A_j \quad \text{and} \quad P_S^m = \textstyle\sum_{j=1}^{Q} P_{Sj}^m = P_S^*$$

For each strategy, the total areas $A^{i*}$ and $A^{m*}$ required to produce the quantity $P_S^*$ under intercropping and sole crop strategies are thus computed as:

$$A^{i*} = \textstyle\sum_{\mathrm{j}=1}^{N} F * A_j \quad \text{and} \quad A^{m*} = \textstyle\sum_{j=1}^{Q} F * A_j \quad \text{where} \quad N \geq Q$$

In the intercropping strategy, maize is simultaneously grown with soybean on the same area $A^{i*}$. Maize productivity on these same $N$ grid-cells, $P_M^i$, as computed as:

$$P_M^i = \sum_{j=1}^{N} P_{Mj}^i = \sum_{j=1}^{N} \bar{Y}_{Mj} * pLER_M * F * A_j$$

with $\bar{Y}_{Mj}$ the average of maize yield projections in the grid-cell $j$ over 2000-2023, $pLER_{\mathrm{M}}$ the pLER of maize, and $P_{Mj}^i$ being the mean productivity of maize in intercropping in the grid-cell $j$.

Finally, in the sole crop strategy, maize is supposed to be grown on the residual area $A^{i*} - A^{m*}$, covered by the $N - Q$ pixels. The production of maize in pure stands is thus estimate as:

$$P_M^i = \sum_{j=1}^{N-Q} P_{Mj}^i = \sum_{j=1}^{N-Q} \bar{Y}_{Mj} * F * A_j$$

### Supplementary Note 3. Training dataset constitution and pre-processing steps of data analyses

#### 1. Crop yields

Soybean and maize yield data were taken from the global dataset of historical yields, which provides worldwide 0.5° (~55 km²) grid-wise data covering the 1981-2016 period (1). Yield values reported in this dataset result from the combination of several sources of information, including national scale yield statistics from the Food and Agriculture Organization, global crop calendars and harvested areas, and satellite-derived net primary production values.

To constitute the soybean and maize yield training dataset of the predictive models, grid-cells from a set of locations representative of EU and global production was constituted. For soybean, data were only available in Italy, so grid-cells located in major soybean producers including Argentina, Brazil, Canada, China, India, and US were additionally included. For maize, grid-cells located in the EU, as well as in Northern China and in the US were included. This choice was motivated by the proximity between maize cropping periods and management in both countries and in the EU. Grid-cells within these locations and with substantial crop area (i.e., where soybean and maize harvested area individually occupied at least 1% of the total cropland) were selected. For each grid-cell, fractions of harvested areas of each crop were taken from the M3 crop mask (2).

To avoid any confusion with technological progress due to improved cultivars and technological progress, yield data were detrended. For a given site and a given crop, yield time series are used to fit a cubic smoothing spline *f(t)* (3, 4). Yield for each year t of this site is then expressed as:

$$Detrended\ yield(t) = f(tmax) + A(t)$$

with *f(tmax)*, defined as the expected yield value at the most recent year of available data for this site (generally 2015 or 2016), and *A(t)*, the yield anomaly, calculated as the difference between expected yield *f(t)* and actual yield in year *t*. This allowed to remove the systematic temporal component of yield that is attributed to technological progress (e.g., genetic improvement, management, inputs). What remains is the interannual variability around that trend, which is typically interpreted as being driven by climate, weather, soil, or other non-progress-related factors.

Previous work showed that increasing the range of environmental conditions in the training dataset improves the accuracy of predictive models (5). Following the procedure employed in previous work (3, 4), several grid-cells located in areas characterized by climate deemed inappropriate for soybean and maize cultivation and resulting to zero yields (such as deserts and arctic areas) were included. These grid-cells were randomly drawn from six zones identified as environmentally improper for any crop production based on the Köppen-Geiger climate classification (6):

- Arid climate zone characterized by winter dryness and hot arid temperatures, coded as **BWh** in the Köppen-Geiger classification (identification number: 7).
- Arid climate zone characterized by winter dryness and cold arid temperatures, coded as **BWk** (identification number: 8).
- Snow climate zone characterized by fully humid conditions, cool summers, and cold winters, coded as **Dfc** (identification number: 20).
- Snow climate zone characterized by fully humid conditions and an extremely continental climate, coded as **Dfd** (identification number: 21).
- Polar climate zone characterized by polar frost conditions, coded as **EF** (identification number: 30).

- Polar climate zone characterized by tundra conditions, coded as **ET** (identification number: 31).

The geographical distribution of these zone is available at: http://koeppen-geiger.vu-wien.ac.at/present.htm.

The selection process was designed to ensure a balanced distribution of sites across climate zones, with the objective of including 20% of zero yield values in the global yield dataset.

### 2. Irrigation fraction

Using the 0.08°-gridded SPAM dataset (7), the fractional area of irrigated soybean (i.e., the proportion of soybean cultivation under irrigation) was retrieved for each grid-cell. A fractional area of 0 indicates that 100% of soybean grown within the considered grid-cell is rainfed. The SPAM data are currently available globally for the year 2020 (v2.0). To ensure consistency with yield data, irrigation data were resampled to a spatial resolution of 0.5° using the *aggregate()* function of the terra R package.

### 3. Climate data

Climate predictors were derived from ERA5-land dataset, a product of the European Centre for Medium-Range Weather Forecasts atmospheric reanalysis of the global climate (8). This dataset provides hourly and monthly estimates of a numerous climate variables at a spatial resolution of 0.1° since January 1950. This dataset is publicly accessible through the Climate Data Store (CDS).

Using the application programming interface of CDS, we directly obtained climate data aggregated at daily frequency from 1981 to 2016 (https://cds.climate.copernicus.eu/cdsapp#!/dataset/reanalysis-era5-land) and resampled at a 0.5° resolution to align with the yield dataset. A set of 10 climate features including minimum and maximum temperatures at two meters, minimum and maximum dewpoint temperatures at two meters (all in K), average precipitation (m), surface net solar radiation (i.e., the difference between downward and reflected solar radiation, J $m^{-2}$), U and V components of wind speed at 10 meters (m $s^{-1}$), and surface pressure (Pa) was retrieved.

Subsequently, six climate variables were derived from the initial variables:

- **Maximum and minimum temperatures (in °C)**: Temperature of air at 2 m above the surface of land, sea, or inland waters. Preprocessing involved the conversion of data(initially expressed in K) to °C. The Copernicus Climate Data Store variables used are: *max_2m_temperature* and *min_2m_temperature*.
- **Average precipitation (in mm)**: Average precipitation height, including liquid and frozen water such as rain and snow, falling on the Earth's surface. Preprocessing involved the conversion of data (initially expressed in m) to mm. The Copernicus Climate Data Store variable used is: *total_precipitation*.
- **Surface net solar radiation (in MJ·$m^{-2}$)**: Amount of solar radiation reaching a horizontal surface at the Earth's surface, including both direct and diffuse radiation, minus the amount reflected by the Earth's surface. Preprocessing involved the conversion of data (initially expressed in J·$m^{-2}$) to MJ·$m^{-2}$. The Copernicus Climate Data Store variable used is: *surface_net_solar_radiation*.
- **Vapor pressure deficit** (in kPa): Difference between the amount of moisture in the air and the amount of moisture the air can hold when saturated. Preprocessing involved calculating the

deficit from mean 2 m air temperature and mean 2 m dewpoint temperature. The Copernicus Climate Data Store variables used are: *2m_temperature (min and max)* and *2m_dewpoint_temperature (min and max).*

- **Reference evapotranspiration (in mm·day$^{-1}$)**: Evapotranspiration rate from a reference surface, representing a hypothetical grass crop that is not short of water. Preprocessing involved calculation using the Penman–Monteith method based on radiation, air temperature, humidity, and wind speed. The Copernicus Climate Data Store variables used are: *2m_temperature (min and max), 2m_dewpoint_temperature (min and max), 10m_u_component_of_wind, 10m_v_component_of_wind, surface_net_solar_radiation,* and *surface_pressure*.

### 4. Aggregation of climates predictors

Based on previous work (3), daily climate predictors were aggregated following three different approaches.

#### *4.1. Monthly averages*

For each site-year, monthly averages of each climate predictor were computed over soybean and maize growing season. For each crop, growing season was delineated on a country-specific basis according to crop calendars sourced from the Agricultural Market Information System.

For soybean (https://www.amis-outlook.org/amis-about/calendars/soybeancal/en/), growing season spanned seven months ranging from:

- November 1st to the next May 31st for sites in Argentina and Brazil,
- from May 1st to November 30th in sites located in Canada,
- from June 1st to December 31st in sites located in India,
- from April 1st to October 31st in sites located in European Union, United-States, and China.

For maize (https://www.amis-outlook.org/amis-about/calendars/maizecal/en/), growing season spanned eight months ranging from April 1st to November 30th in all sites, which were selected in the Northern hemisphere (including European Union, United-States, Northern China).

This led to a total of 42 (seven months * six climate variables) and 48 (eight months * six climate variables) monthly climate predictors for soybean and maize datasets, respectively.

#### *4.2. Seasonal averages*

For each site-year and each crop, the mean of each climate variable over the growing season was computed. This technique reduced the number of predictors to six (i.e., one per climate variable).

#### *4.3. Principal component analysis*

The last approach used was principal component analysis (PCA) (9). The central concept of PCA is to compute new variables called principal components (PC) from linear combinations of the original predictors. The first principal component is required to have the largest possible variance and therefore this component will explain the largest part of the variance in the predictor dataset. The second component is computed under the constraint of being orthogonal to the first component and to have the largest possible explained variance.

For all climate variables, the first three PCs, hereafter referred to as PC1, PC2, and PC3, explained more than 90% of the variability in monthly climate data (Supplementary Figure 25).

PC1 of each variable was highly correlated to all monthly averages, with similar direction and generally similar strength across months, while PC2 gave more weights to some specific months (left panels of Supplementary Figures 5 to 8). This suggests that PC1 summarized the average climate conditions during the growing season, while PC2 gave more focus on specific periods.

### Supplementary Note 4. Assessment of the predictive performance of several crop yield forecasting models

#### 1. Tested models

In this study, a random-forest (RF) model was used to predict crop yield. RF (10) is a tree-based machine-learning method that makes no assumption regarding the distribution and relationship between predictors and yield. Briefly, it consists in building an ensemble of independent decision trees from bootstrapped samples. Individual trees have the properties to have low bias but high variance, and when combined together, produce an output with lower variance. RF was chosen because this machine learning algorithm showed good performance compared to other algorithms in predicting yield of crops (11), especially soybean (3, 4).

For each crop, four predictive models based different climate predictors aggregation methods were built:

(i) a model based on *monthly averages* of climate variables (corresponding to 42 or 48 climate predictors depending on the crop),
(ii) a model based on *seasonal averages* of climate variables (corresponding to six climate predictors in total),
(iii) a model based on the *two first PCs* of each climate variable (corresponding to 12 climate predictors in total), and
(iv) a model based on the *three first PCs* of each climate variable (corresponding to 12 climate predictors in total). This led to train eight models in total.

#### 2. Predictive performance metrics

The predictive performance of the models was assessed using the Nash-Sutcliffe model efficiency (NSE, unitless) as predictive performance metric. An efficiency of one corresponds to a perfect match of predictions to observed data, an efficiency of zero indicates that predictions are as accurate as the mean of observed data, whereas an efficiency lower than zero occurs when the observed mean is a better predictor than the tested model.

#### 3. Cross-validation strategies

Previous articles emphasized the importance of rigorous cross-validation strategies to ensure that the predictive performance of a given model is evaluated on a dataset independent from the one used to train that algorithm. For each model, NSE was estimated by separate cross-validation strategies to evaluate temporal and spatial extrapolation.

First, a year-by-year cross-validation was performed, to assess model's capability in predicting yields in a new year, not included in the training dataset (temporal extrapolation).

Secondly, a group-wise cross-validation was employed, wherein 10 randomly selected site groups were used to evaluate the model's ability to forecast yields in novel geographic regions not encompassed within the training dataset (spatial extrapolation).

**Supplementary Note 5. Computation of partial land equivalent ratios of soybean and maize for sensitivity analyses including the effect of fertilization and temporal niche differentiation on partial land equivalent ratios**

We used the regression models published in the meta-analysis of Xu et al. (12) to estimate the partial land equivalent ratios (pLERs) in a maize-soybean intercropping system in two different scenarios: 1) assuming a high N fertilization rate; 2) assuming that both species are sown and harvested at the same time, corresponding to a null temporal niche differentiation (TND).

1. **Scenario with high N inputs**

We assume that fertilizer use in maize-soybean intercropping will be equivalent to the one for maize, which is generally more fertilized than soybean. To be representative of local fertilization practices in the European Union (EU), we used the N fertilizer applications rates derived from the NPKGRIDS dataset (11). This dataset reports the average N rate applied on several crops (including maize) in 2020 at a resolution of 0.05°. Data were aggregated at a 0.5° resolution to be consistent with the resolution used in this study.

Xu et al. (12) related the pLER of maize and the pLER of soybean to additional fertilizer N. They report the following regressions:

- pLER of maize = 0.767 + 0.00011 * N (Fig. 8b of the paper; P = 0.6322)
- pLER of soybean = 0.591 – 0.000583 * N (Fig. 8c of the paper; P = 0.0131).

Using these equations and the local rates described above, we obtained the site-specific pLERs for soybean and maize (see Supplementary Figure 10). **These values ranging from 0.45 and 0.59 for soybean and between 0.77 and 0.79 for maize**, respectively.

2. **Scenario with null TND**

In Xu et al. (12), the average land equivalent ratio (LER) provided by the same study was 1.32, with pLERs values of 0.56 and 0.79 for soybean and maize, respectively.

Additionally, the authors report a significant relationship between LER and TND with a slope of 0.214 ± 0.011 (units of LER per unit of TND) and an intercept (LER at TND = 0) of 1.25 ± 0.03.

We computed the ratio between each individual pLER and average LER (i.e., 0.56/1.32 = 0.42 for soybean and 0.79/1.32 = 0.60 for maize) and we used them to weight the LER at TND = 0 to derive corresponding pLERs.

We ended with **pLERs values of 0.53 and 0.76** for soybean and maize, respectively.

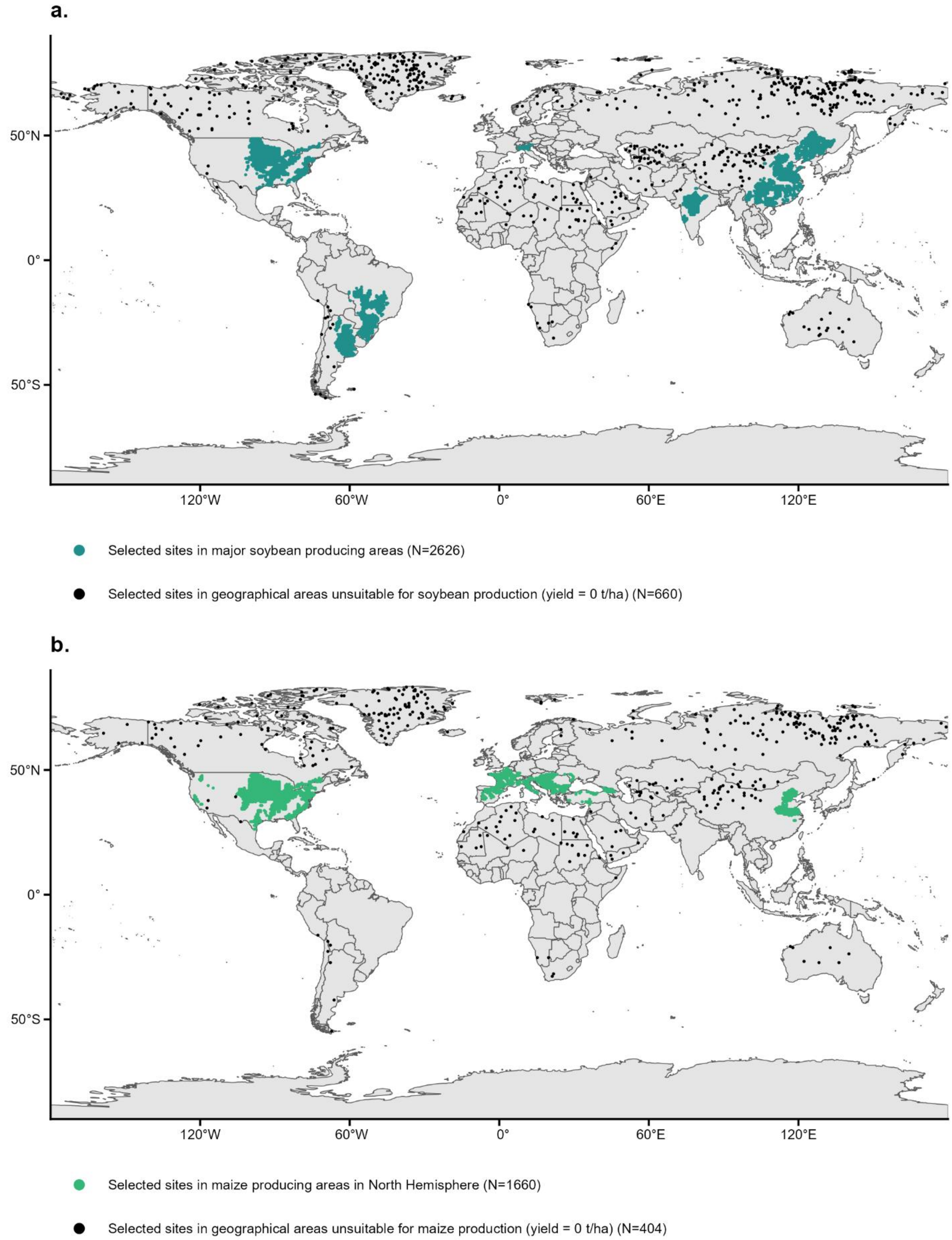


**Supplementary Figure 1. Geographical distribution of sites included in soybean (a) and maize (b) the training dataset.** Soybean and maize training datasets included 116,516 and 72,964 unique combinations of sites and years, respectively. Base map based on Natural Earth data, created using the R package rnaturalearth.

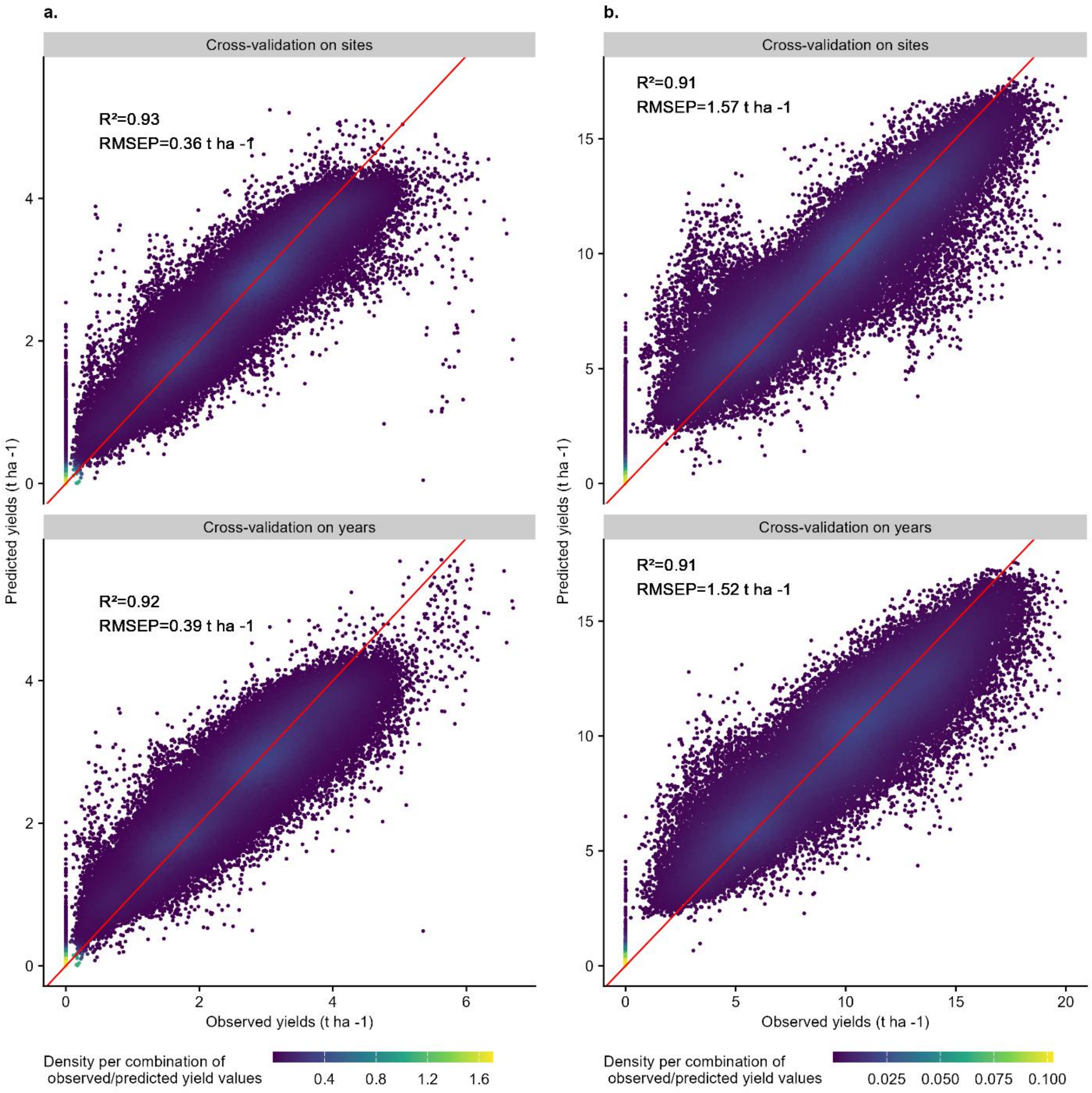


**Supplementary Figure 2. Scatterplots of soybean (a) and maize (b) yields observations compared to the best model predictions estimated by cross-validation on sites (top panel) or on years (bottom panel).**

Best model was the random forest model based on the two first principal components derived from monthly climate data. The red line in scatterplots represents the 1:1 line. Soybean and maize training datasets included 116,516 and 72,964 unique combinations of sites and years, respectively.

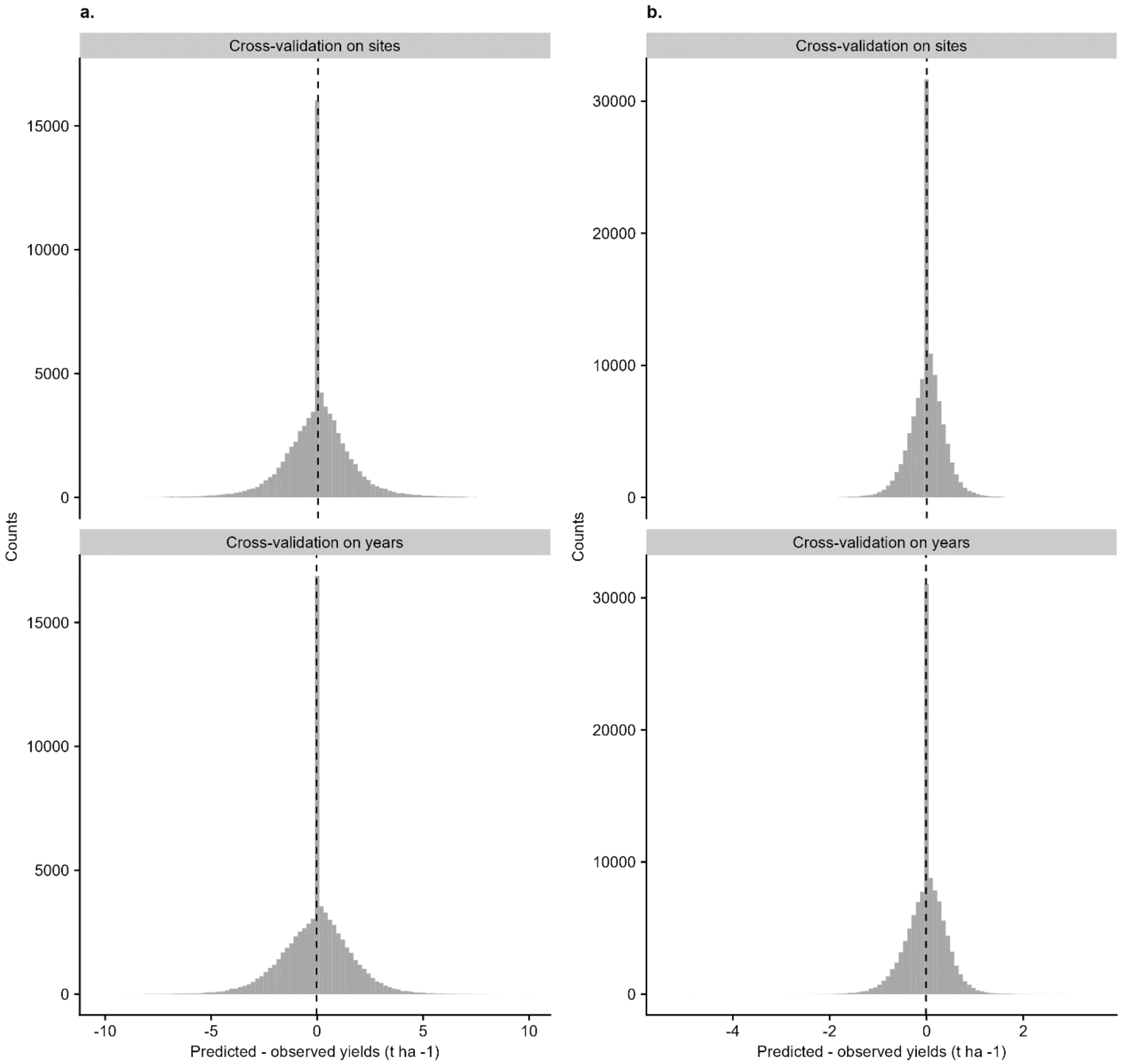


**Supplementary Figure 3. Distribution of the residuals (predictions - observation) of the best models forecasting soybean (a) and maize (b) yields, estimated by cross-validation on sites (top panel) or on years (bottom panel).**

Best model was the random forest model based on the two first principal components derived from monthly climate data. Black line in residues histogram represents mean residuals. Soybean and maize training datasets included 116,516 and 72,964 unique combinations of sites and years, respectively.

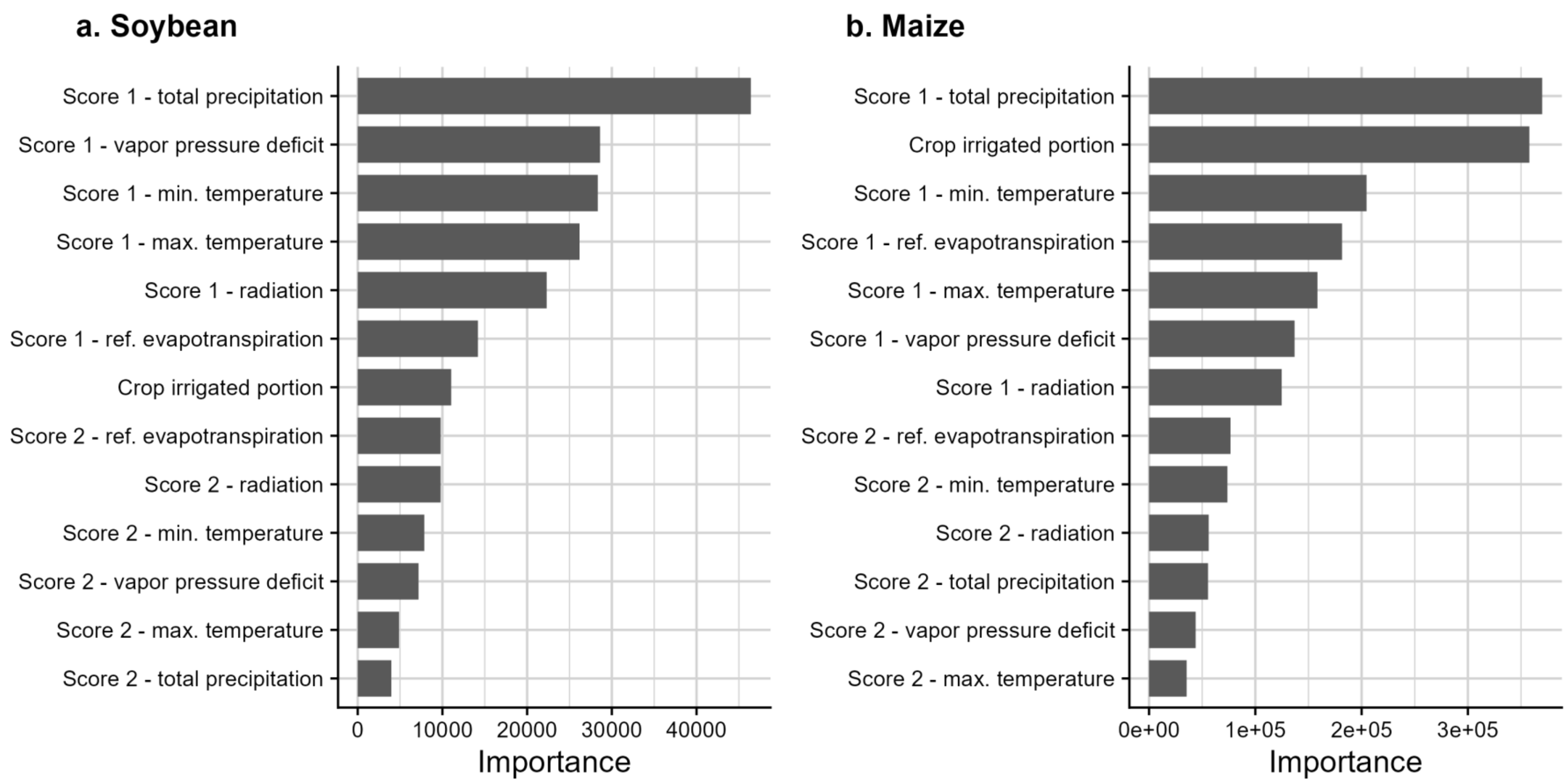


**Supplementary Figure 4. Importance of predictors in the random forest models showing best performance to predict soybean (a) and maize (b) yields in the training datasets.**

Best models were the random forest models based on two principal components ("scores 1", or "score 2") derived from principal component analysis based on monthly climate data, respectively. Soybean and maize training datasets included 116,516 and 72,964 unique combinations of sites and years, respectively.

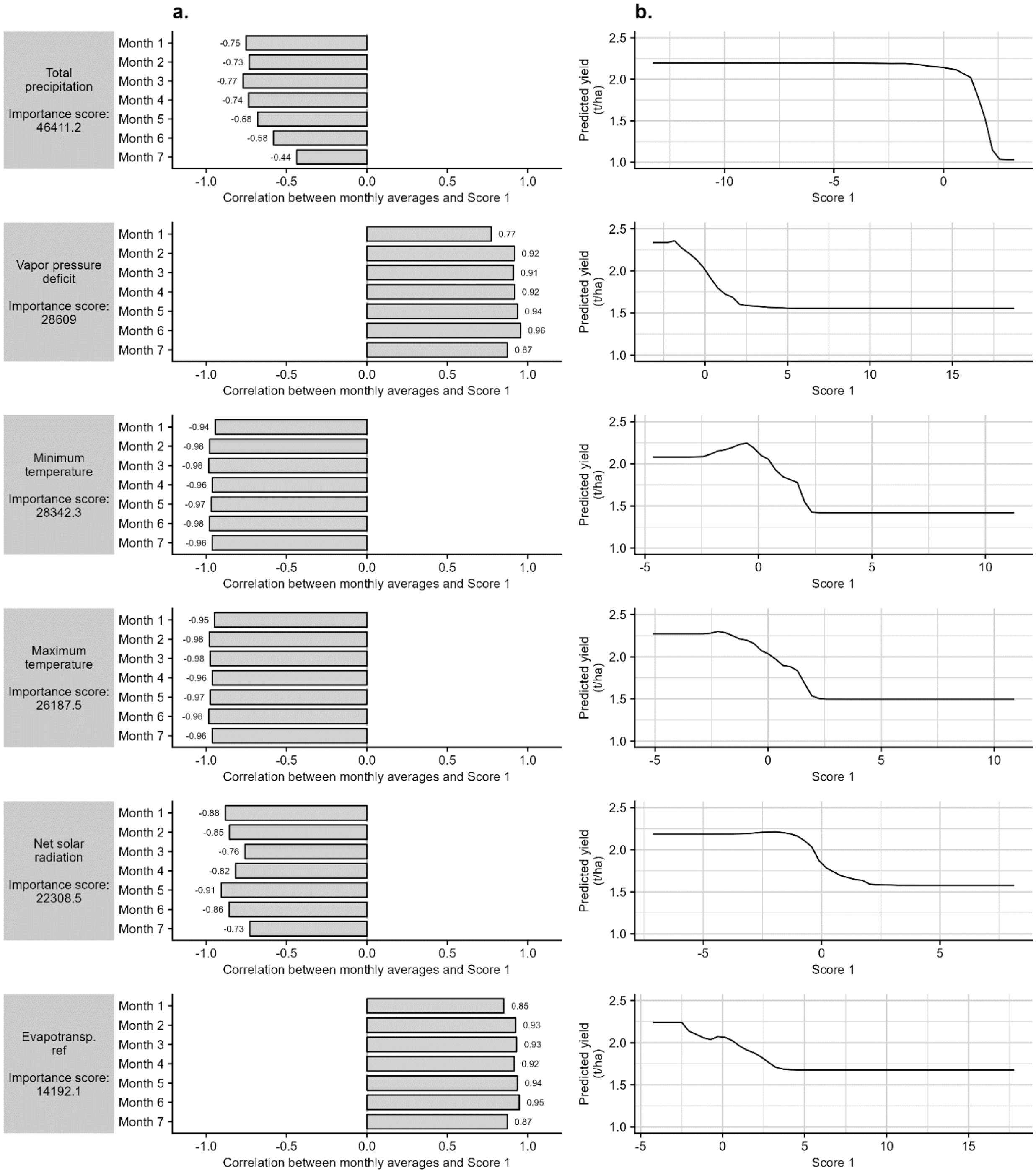


**Supplementary Figure 5. Contribution of climate data to the first principal component derived from principal component analysis applied on monthly averages (a) and partial dependency plot associated with each predictor (b) in the best model predicting *soybean* yield.**

Best model was the random forest model including the scores associated with the first and second components derived from principal component analysis applied on monthly climate data. Climate variables ordered by importance in the model. Panel a: for each climate variable, height and direction of the bars indicate Pearson correlation coefficient between monthly averages and the first principal component (i.e. "score 1"). Panel b: the expected value of model yield prediction is plotted as a function of each predictor. Soybean training dataset included 116,516 unique combinations of sites and years.

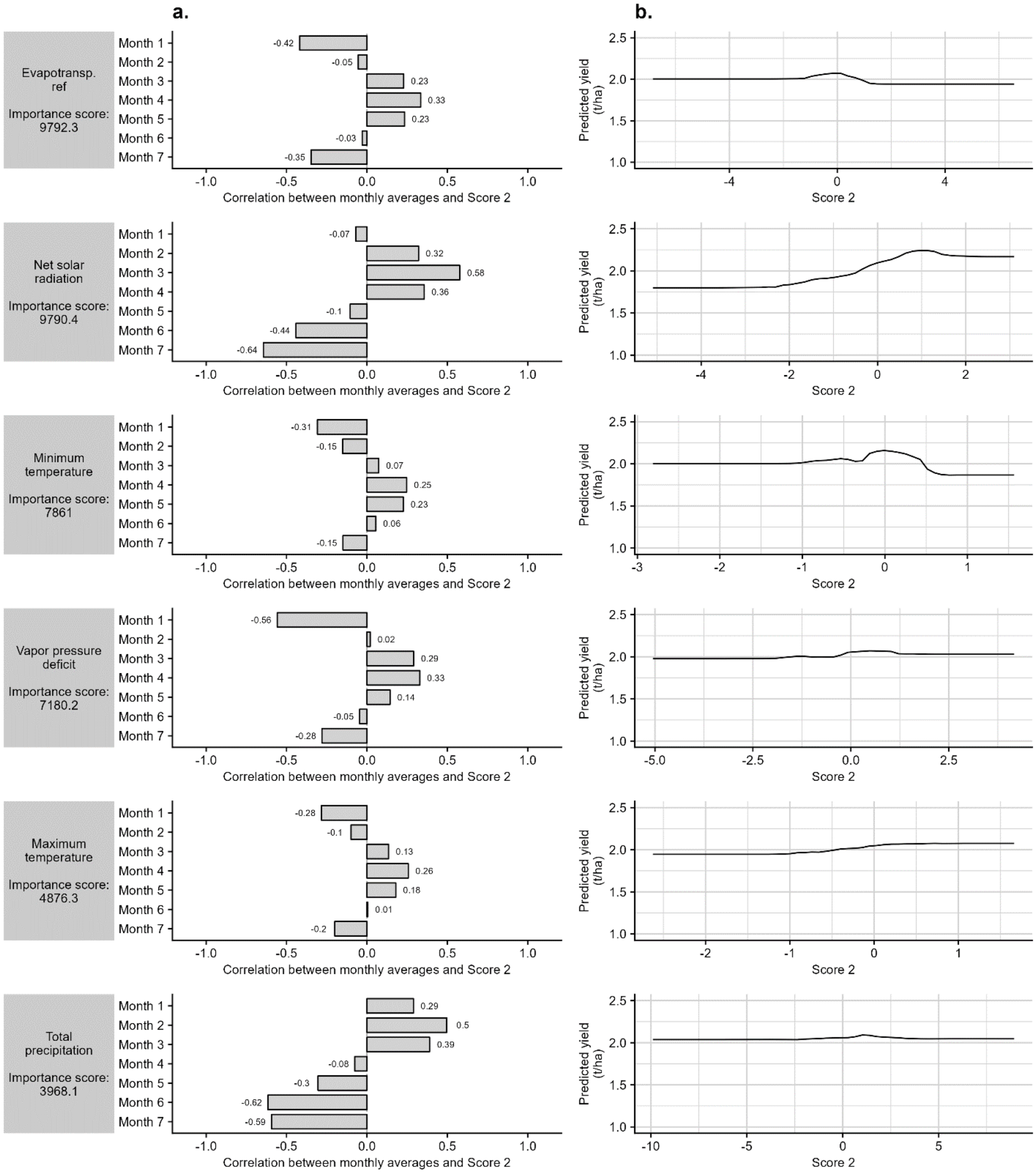


**Supplementary Figure 6. Contribution of climate data to the second principal component derived from principal component analysis applied on monthly averages (a) and partial dependency plot associated with each predictor (b) in the best model predicting *soybean* yield.**

Best model was the random forest model including the scores associated with the first and second components derived from principal component analysis applied on monthly climate data. Climate variables ordered by importance in the model. Panel a: for each climate variable, height and direction of the bars indicate Pearson correlation coefficient between monthly averages and the second principal component (i.e. “score 2”). Panel b: the expected value of model yield prediction is plotted as a function of each predictor. Soybean training dataset included 116,516 unique combinations of sites and years.

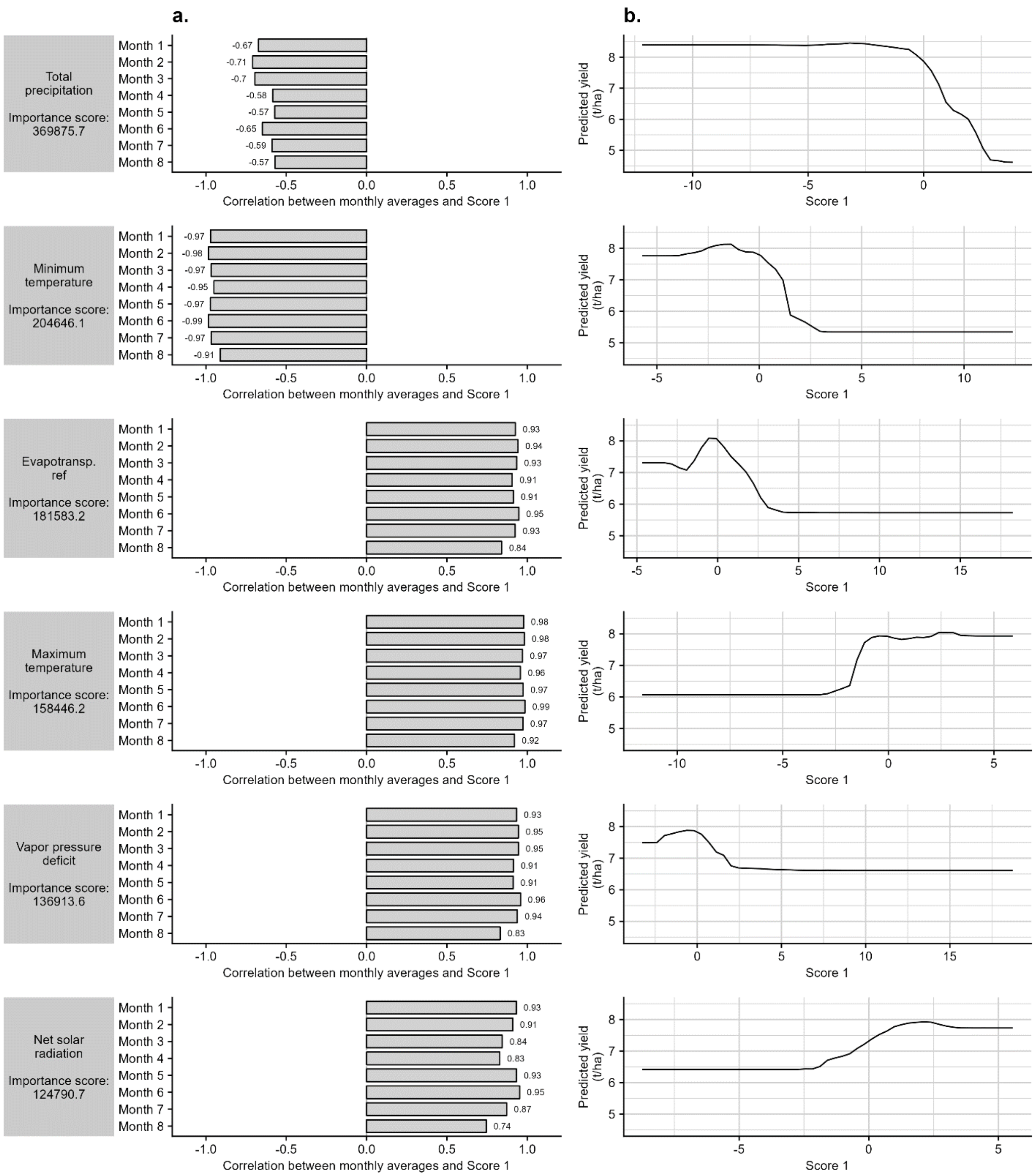


**Supplementary Figure 7. Contribution of climate data to the first principal component derived from principal component analysis applied on monthly averages (a) and partial dependency plot associated with each predictor (b) in the best model predicting *maize* yield.**

Best model was the random forest model including the scores associated with the first and second components derived from principal component analysis applied on monthly climate data. Climate variables ordered by importance in the model. Panel a: for each climate variable, height and direction of the bars indicate Pearson correlation coefficient between monthly averages and the first principal component (i.e. "score 1"). Panel b: the expected value of model yield prediction is plotted as a function of each predictor. Maize training dataset included 72,964 unique combinations of sites and years.

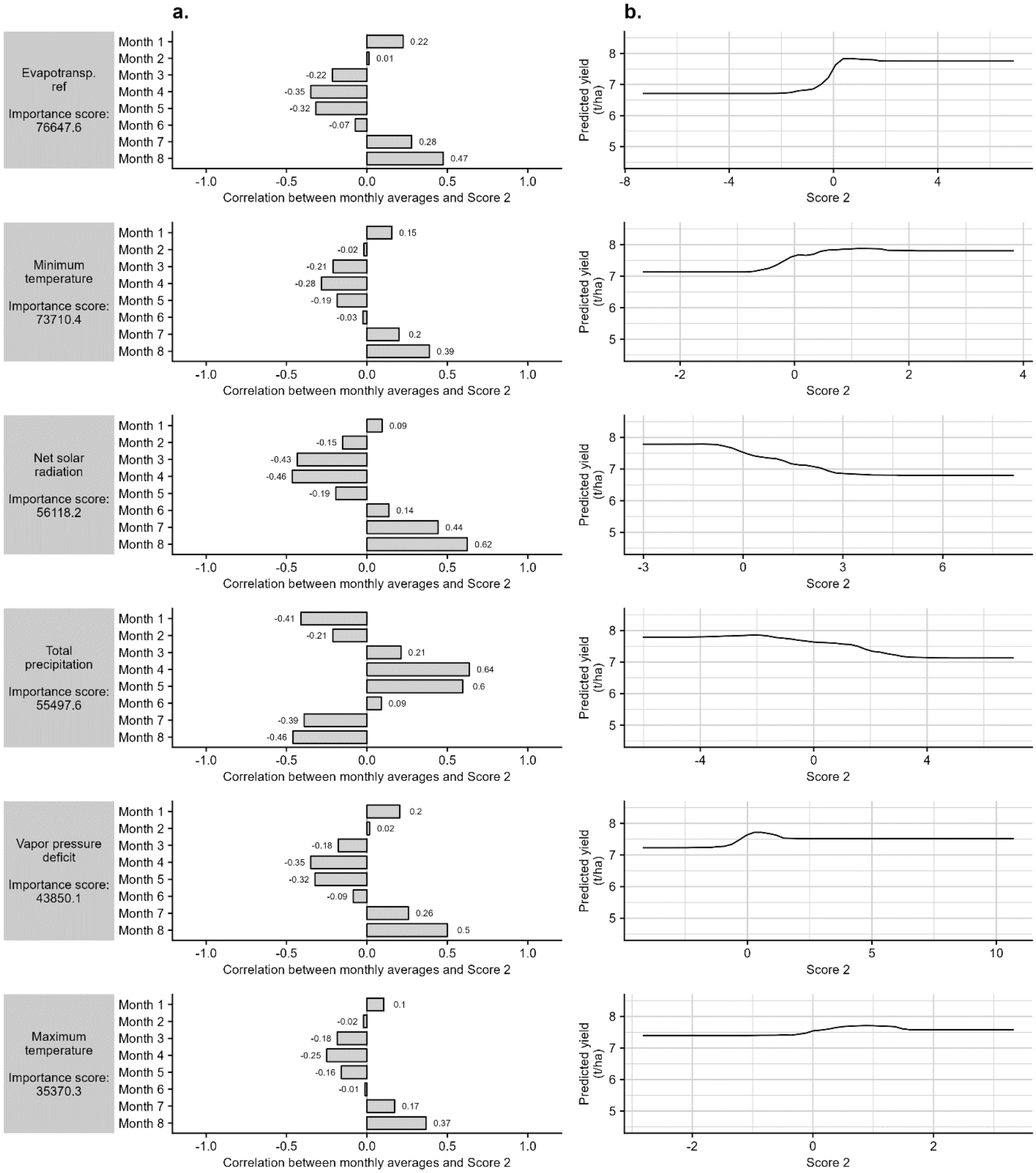


**Supplementary Figure 8. Contribution of climate data to the second principal component derived from principal component analysis applied on monthly averages (a) and partial dependency plot associated with each predictor (b) in the best model predicting *maize* yield.**

Best model was the random forest model including the scores associated with the first and second components derived from principal component analysis applied on monthly climate data. Climate variables ordered by importance in the model. Panel a: for each climate variable, height and direction of the bars indicate Pearson correlation coefficient between monthly averages and the second principal component (i.e. "score 2"). Panel b: the expected value of model yield prediction is plotted as a function of each predictor. Maize training dataset included 72,964 unique combinations of sites and years.

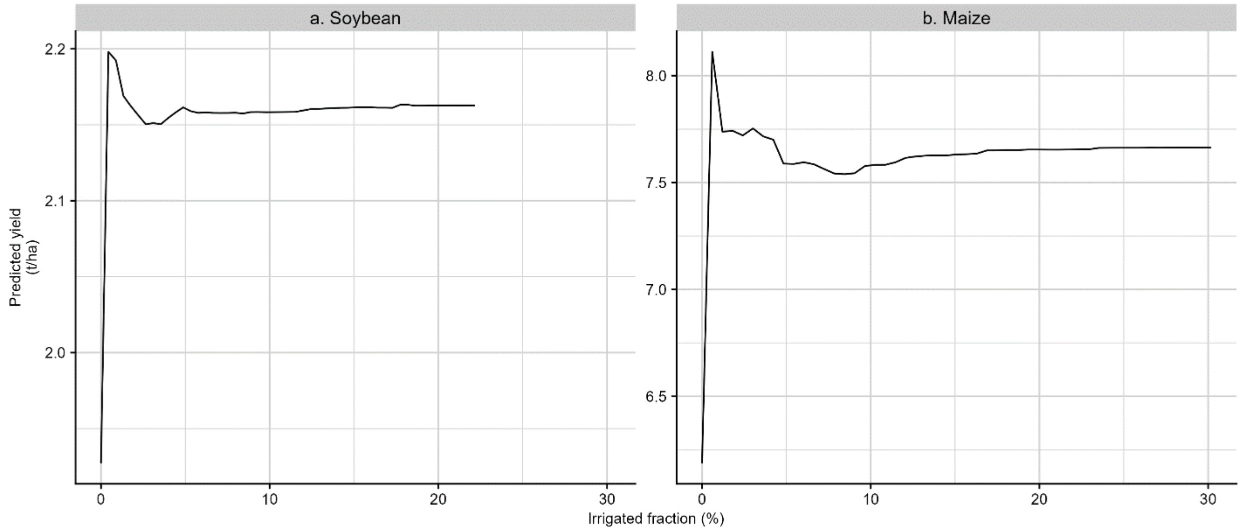


**Supplementary Figure 9. Partial dependency plot associated with irrigated fraction in the best model predicting soybean (a) and maize (b) yield.**

For both crop, best model was the random forest model including the scores associated with the first and second components derived from principal component analysis applied on monthly climate data. Soybean training dataset included 116,516 unique combinations of sites and years. Maize training dataset included 72,964 unique combinations of sites and years.

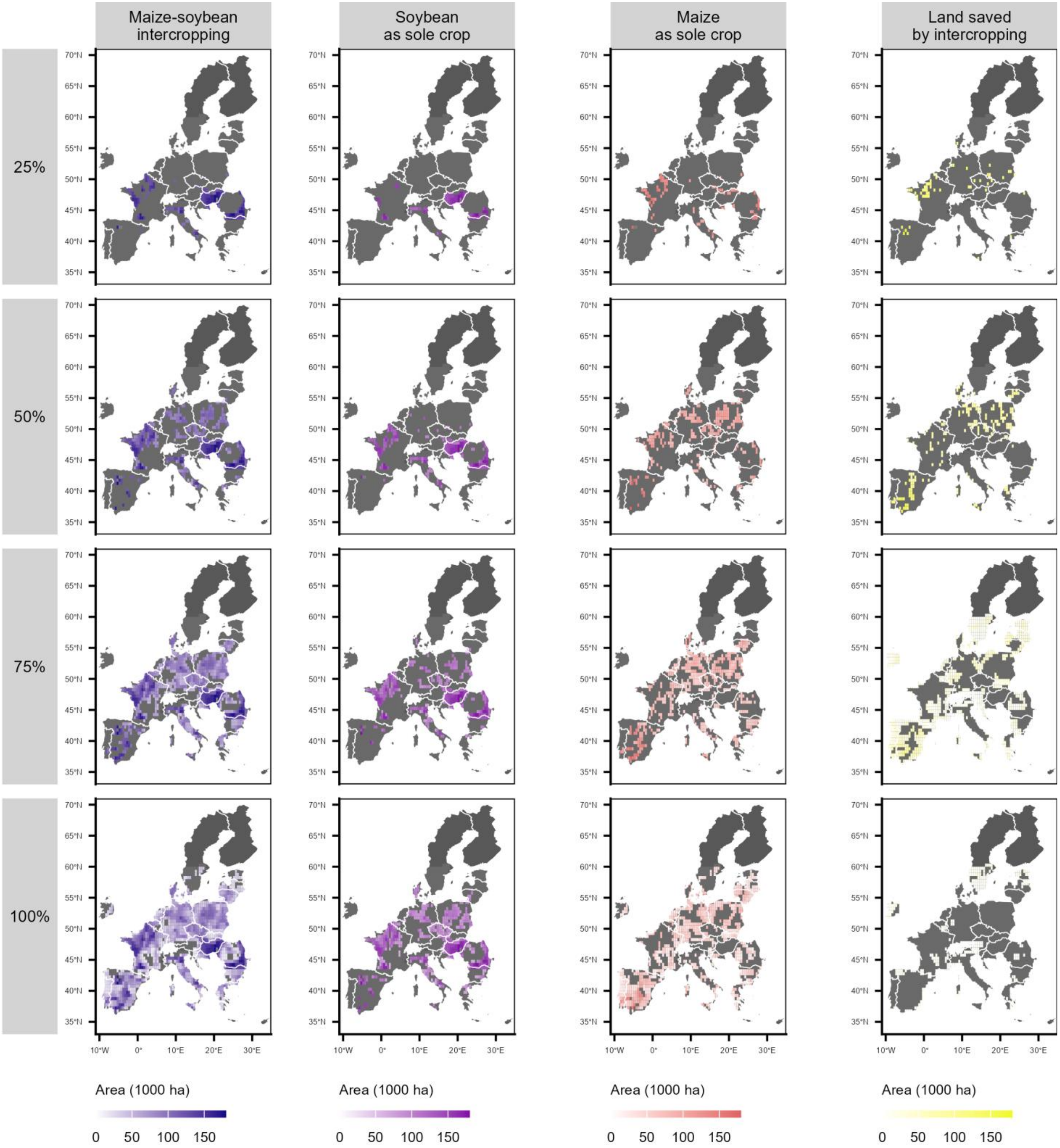


**Supplementary Figure 10. Maize-soybean intercropping and sole cropping allocations for 25, 50, 75, and 100% soybean self-sufficiency (i.e., 9.1, 18.2, 27.2, and 36.3 Mt, respectively) in the European Union.**

First column: Intercropping was sequentially allocated in highest yielding sites, ranked by mean average rainfed soybean production over 2000-2023 period until the considered level of soybean self-sufficiency is met. Second column: sole soybean allocated following the same procedure. Third column: sole maize allocated to the residual area so that the total covered area is equal in both strategies. Fourth column: additional area required to reach the same coproduction in both strategies. All simulations based on (i) rainfed yield projections from a random-forest model based on the two first principal components of climate variables and irrigation fraction, (ii) crop return frequency of one-in-four year, and (iii) partial land equivalent ratios equal to 0.56 and 0.79 for soybean and maize, respectively (12). The maximum total allocated area was set at 25% of croplands in the European Union. Base map based on Natural Earth data, created using the R package rnaturalearth.

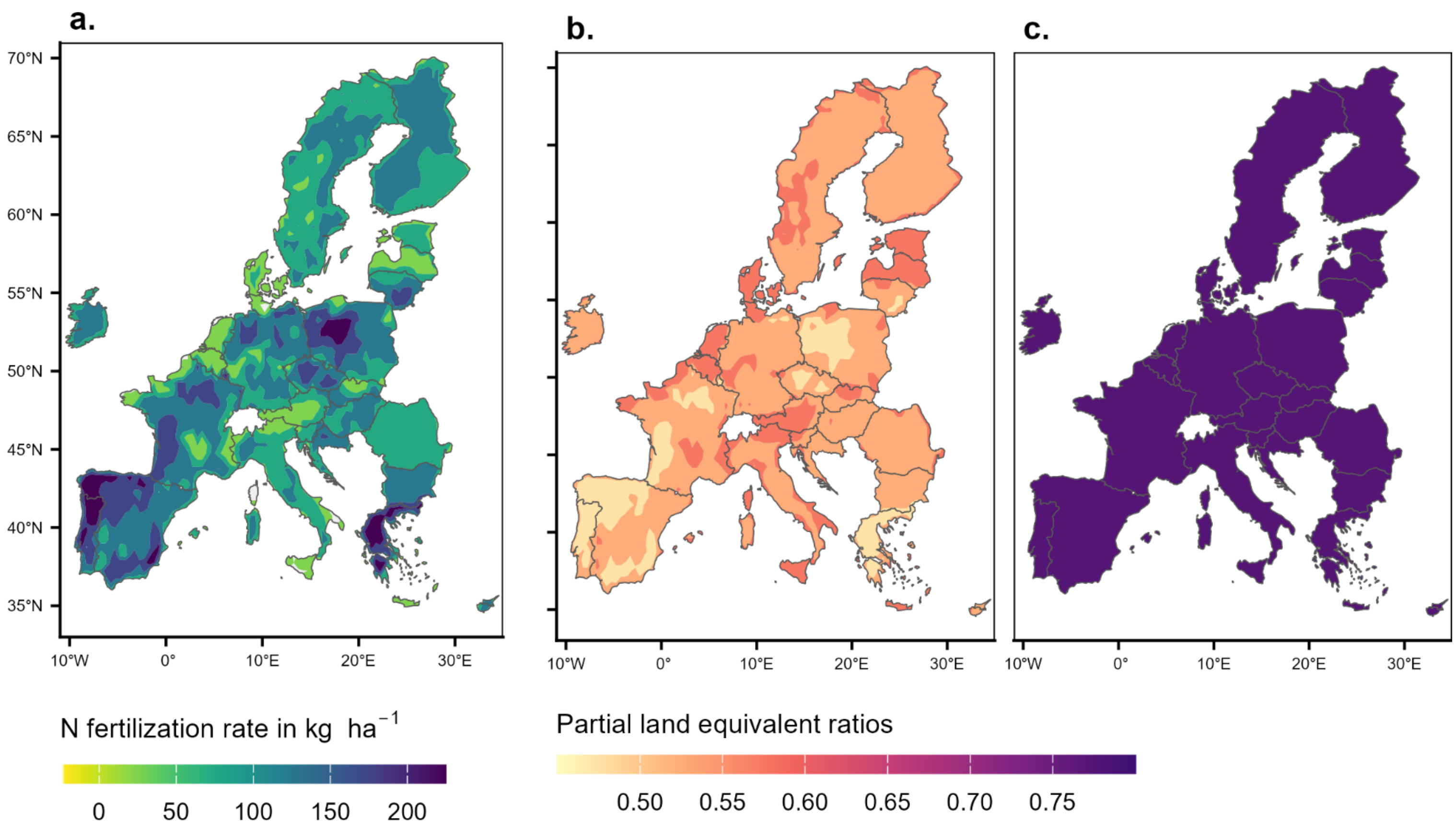


**Supplementary Figure 11. The performances of maize-soybean intercropping according to (a) nitrogen fertilization rates for maize of 2020, represented by the partial land equivalent ratios of (b) soybean and (c) maize.** Gridded fertilization rates (0.5°-resolution) were obtained from the NPKGRIDS dataset (11). Soybean and maize pLERs values were computed based on the equations of Xu et al. (12)**.** Base map based on Natural Earth data, created using the R package rnaturalearth.

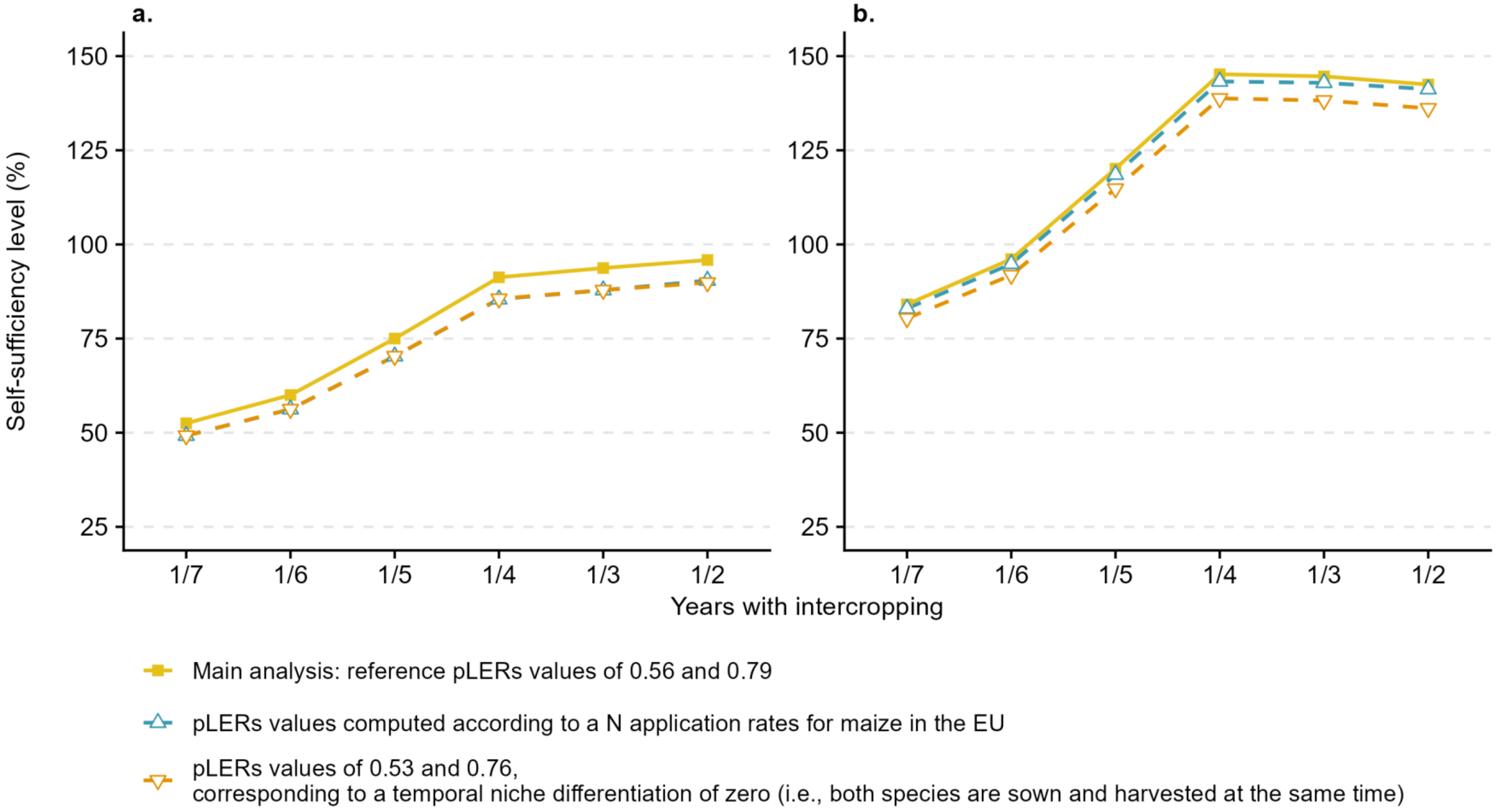


**Supplementary Figure 12. Levels of soybean (a) and maize (b) self-sufficiency in the European Union (EU) achieved from intercropping for different assumptions of partial land equivalent ratios (pLERs) and crop return frequency.**
Crop frequencies of 1/7, 1/6, 1/5, 1/4, 1/3, and 1/2 correspond to allocating maize-soybean intercropping on 14%, 16%, 20%, 25%, 33%, or 50% of cropland area in each grid-cell. Intercropping was first allocated to soybean highest-yielding grid-cells, and in all cases total intercropping area was constrained to not exceed 25% of croplands in the EU (i.e., 25 Mha). Squares represent the results obtained with pLER values of 0.56 for soybean and 0.79 for maize, corresponding to the average productive performances of maize-soybean intercropping systems estimated in a previous meta-analysis (12). The facing down triangles represent the results obtained with site-specific pLER values recomputed according to local N fertilization rate of maize in the European-Union (data from the NPKGRIDS dataset [11]), based on the results of Xu et al. (12). Facing upwards triangles represent the results obtained with pLER values corresponding to a temporal differentiation niche of 0 (i.e., 0.53 for soybean and 0.76 for maize), based on the results of Xu et al. [10]).

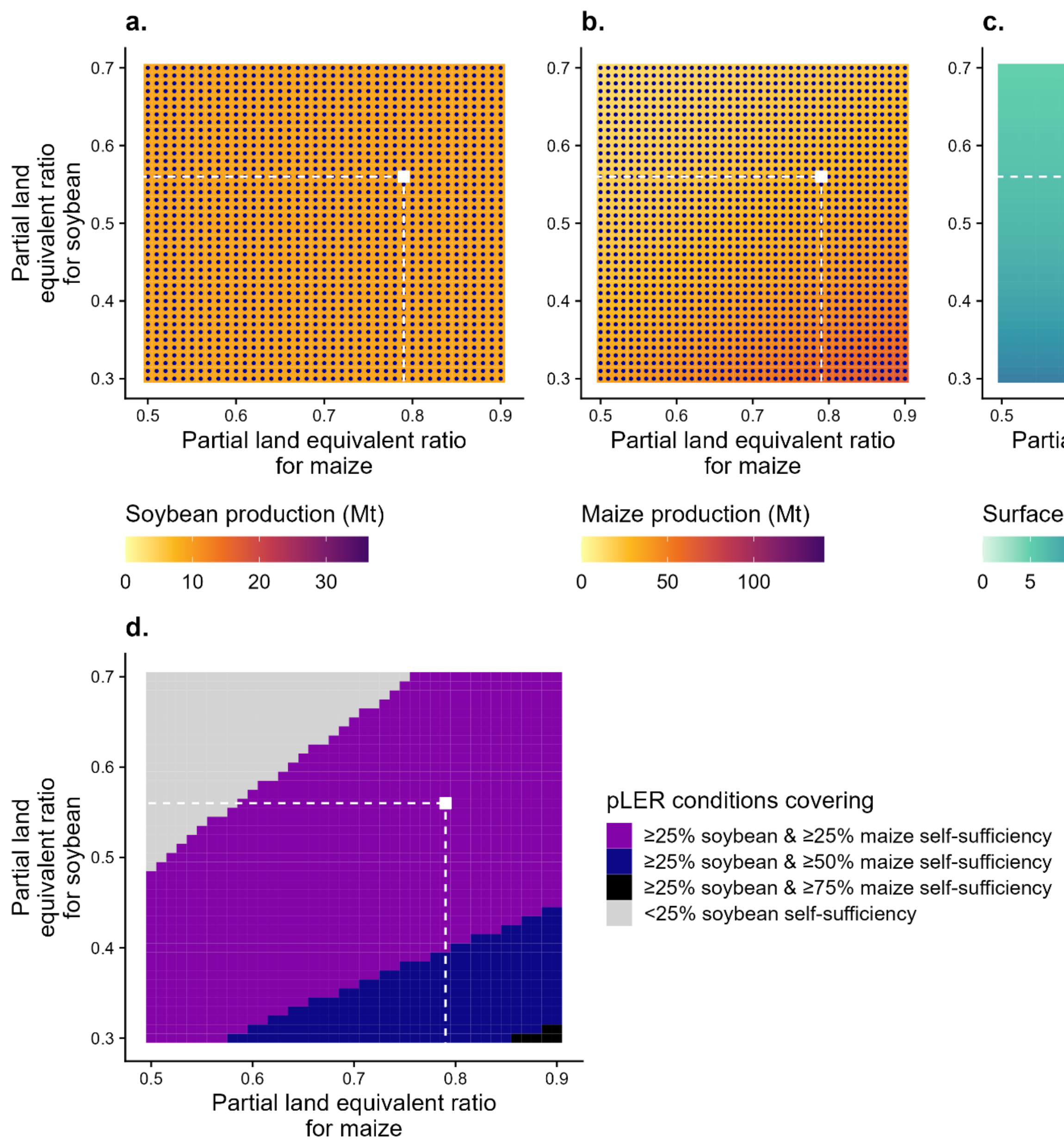


**Supplementary Figure 13. Characteristics of maize-soybean intercropping allocations in terms of soybean produced (a), maize produced (b), and surface allocated (c) for covering 25% soybean self-sufficiency in the European Union depending on intercropping efficiency (pLER). The minimum efficiency conditions simultaneously satisfying 25% of soybean and 25, 50, 75, or 100% maize self-sufficiency are also shown (d).**
Scenarios with soybean and maize productions satisfying 50% and 100% self-sufficiency (i.e., 9.0 and 85.1 Mt, respectively) are highlighted by blue dots in panels (a) and (b), respectively. All scenarios assume a one-in-four year return frequency of intercropping and the total surface is limited to 25% of cropland area in the European Union (i.e., 25 Mha). Scenarios where the surface allocated to intercropping reaches that threshold are represented by the red dots in panel (c). In all panel, the white dot represent the current estimation of maize-intercropping efficiency at a global scale (12).

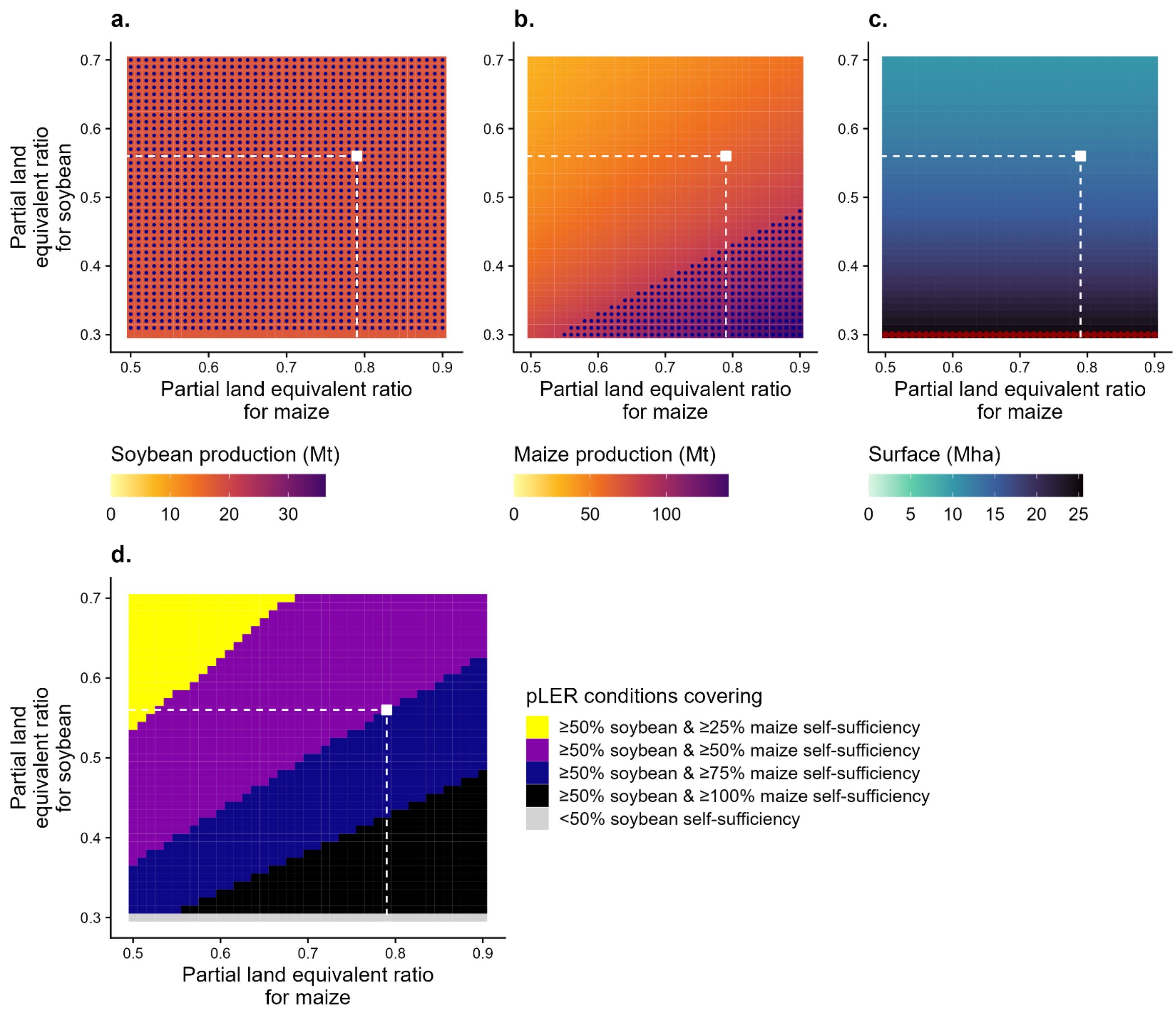


**Supplementary Figure 14. Characteristics of maize-soybean intercropping allocations in terms of soybean produced (a), maize produced (b), and surface allocated (c) for covering 50% soybean self-sufficiency in the European Union depending on intercropping efficiency (pLER). The minimum efficiency conditions simultaneously satisfying 50% of soybean and 25, 50, 75, or 100% maize self-sufficiency are also shown (d).**
Scenarios with soybean and maize productions satisfying 50% and 100% self-sufficiency (i.e., 18.1 and 85.1 Mt, respectively) are highlighted by blue dots in panels (a) and (b), respectively. All scenarios assume a one-in-four year return frequency of intercropping and the total surface is limited to 25% of cropland area in the European Union (i.e., 25 Mha). Scenarios where the surface allocated to intercropping reaches that threshold are represented by the red dots in panel (c). In all panel, the white dot represent the current estimation of maize-intercropping efficiency at a global scale (12).

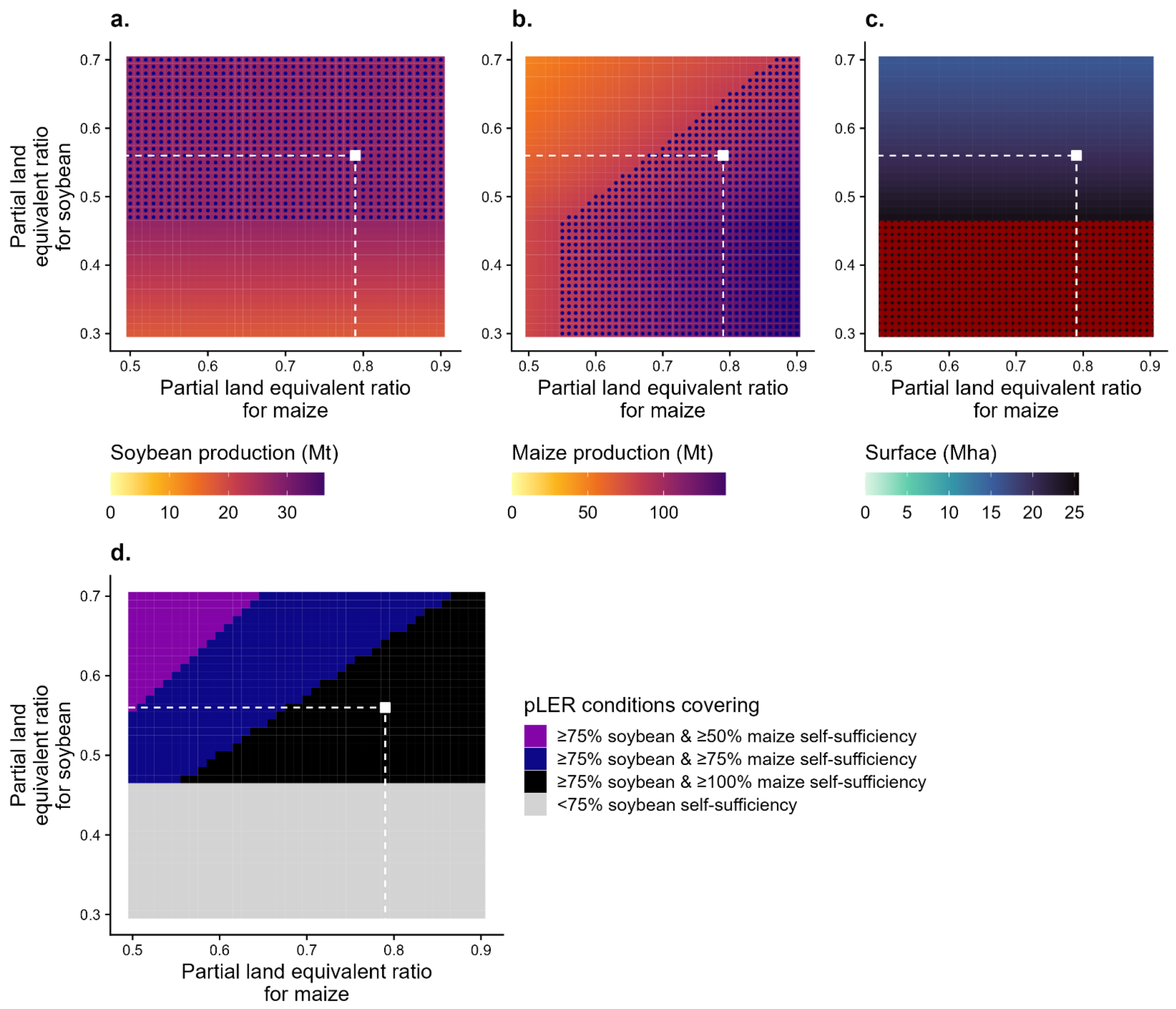


**Supplementary Figure 15. Characteristics of maize-soybean intercropping allocations in terms of soybean produced (a), maize produced (b), and surface allocated (c) for covering 75% soybean self-sufficiency in the European Union depending on intercropping efficiency (pLER). The minimum efficiency conditions simultaneously satisfying 75% of soybean and 25, 50, 75, or 100% maize self-sufficiency are also shown (d).**

Scenarios with soybean and maize productions satisfying 50% and 100% self-sufficiency (i.e., 27.2 and 85.1 Mt, respectively) are highlighted by blue dots in panels (a) and (b), respectively. All scenarios assume a one-in-four year return frequency of intercropping and the total surface is limited to 25% of cropland area in the European Union (i.e., 25 Mha). Scenarios where the surface allocated to intercropping reaches that threshold are represented by the red dots in panel (c). In all panel, the white dot represent the current estimation of maize-intercropping efficiency at a global scale (12).

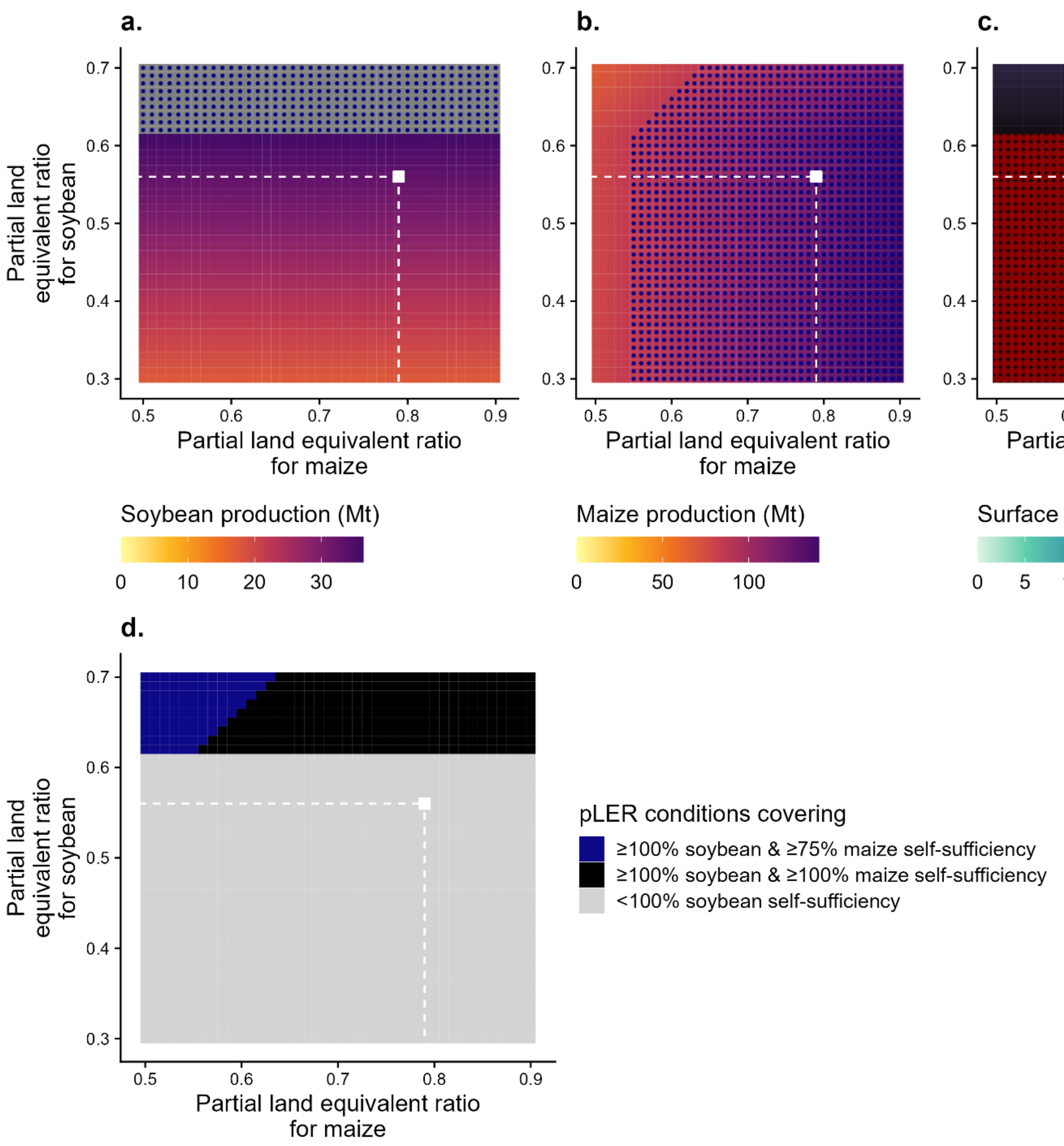


**Supplementary Figure 16. Characteristics of maize-soybean intercropping allocations in terms of soybean produced (a), maize produced (b), and surface allocated (c) for covering 100% soybean self-sufficiency in the European Union depending on intercropping efficiency (pLER). The minimum efficiency conditions simultaneously satisfying 100% of soybean and 25, 50, 75, or 100% maize self-sufficiency are also shown (d).**
Scenarios with soybean and maize productions satisfying 100% and 100% self-sufficiency (i.e., 36.6 and 85.1 Mt, respectively) are highlighted by blue dots in panels (a) and (b), respectively. All scenarios assume a one-in-four year return frequency of intercropping and the total surface is limited to 25% of cropland area in the European Union (i.e., 25 Mha). Scenarios where the surface allocated to intercropping reaches that threshold are represented by the red dots in panel (c). In all panel, the white dot represent the current estimation of maize-intercropping efficiency at a global scale (12).

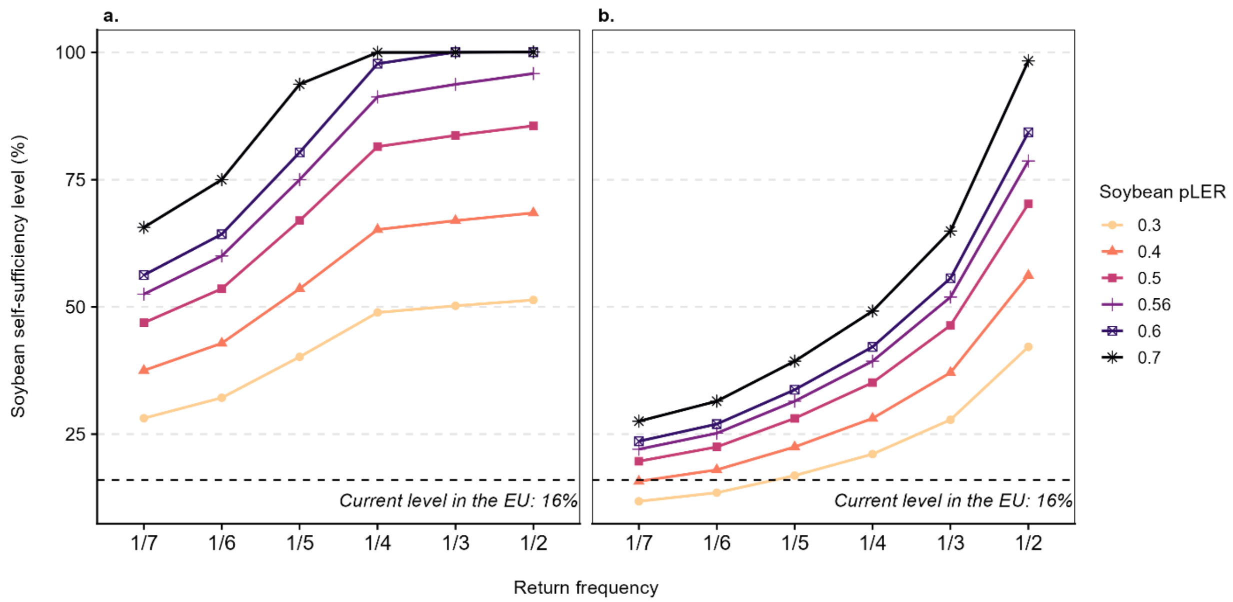


**Supplementary Figure 17. Level of soybean self-sufficiency in the European Union (EU) achieved by intercropping on areas showing soybean productivity equal or higher than 1 $t.ha^{-1}$ (left panel) or 2.6 $t.ha^{-1}$ (right panel), in several scenarios of partial land equivalent ratio (pLER) and crop return frequency.**

Crop frequencies of one year in seven, six, five, four, three, or two correspond to allocating maize-soybean intercropping on 14, 16, 20, 25, 33, or 50% of cropland area in each site. For all scenarios, total cropping area did not exceed 25% of croplands in the EU (i.e., 25 Mha). All simulations are based on (i) yield projections from a random-forest model based on the two first principal components of climate variables and irrigation fraction, (ii) crop return frequency of one-in-four years, and (iii) partial land equivalent ratios equal to 0.56 and 0.79 for soybean and maize, respectively (12).

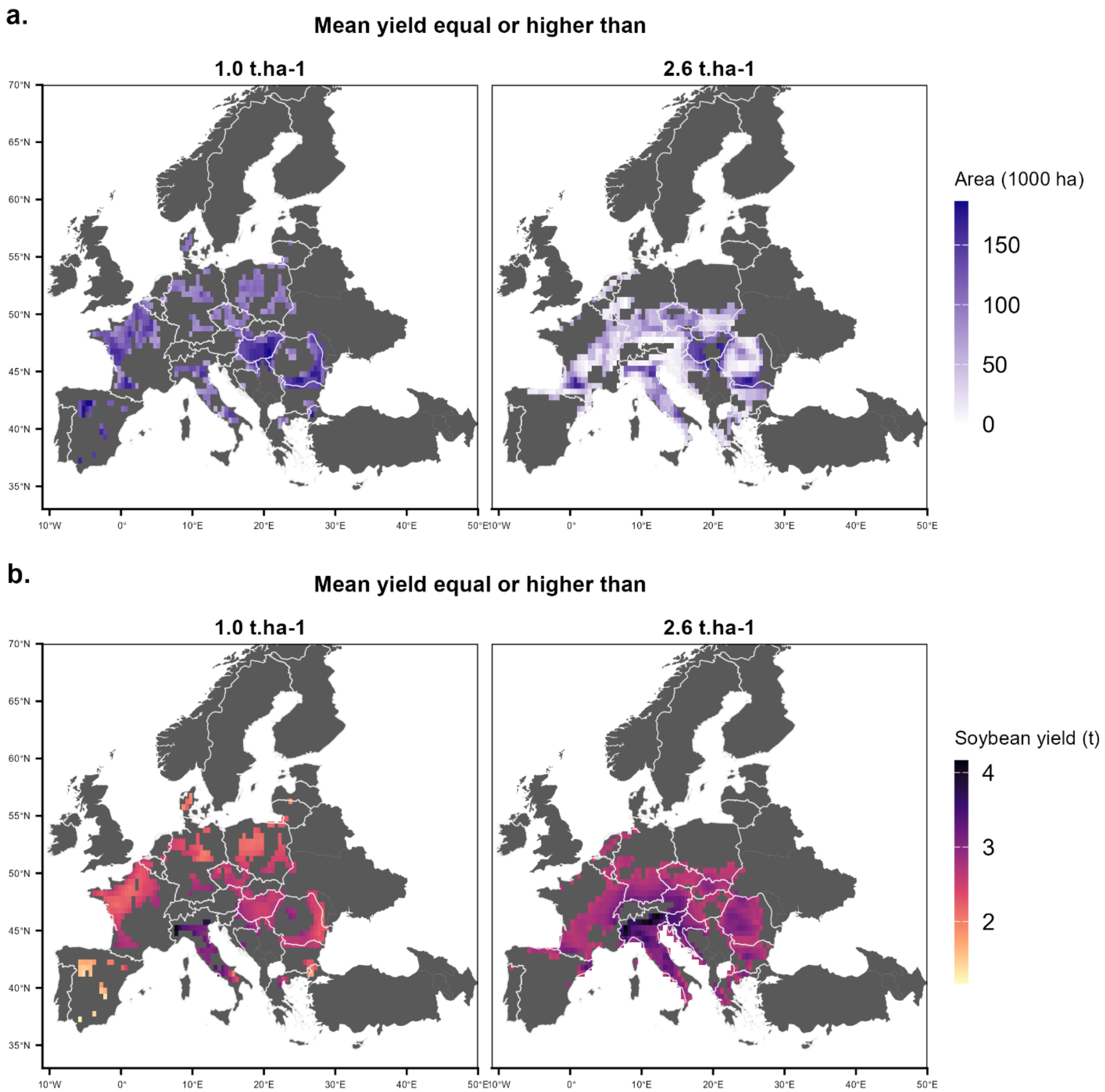


**Supplementary Figure 18. Geographical allocation of maize-soybean intercropping in areas showing soybean productivity equal or higher than 1 t.ha$^{-1}$ or 2.6 t.ha$^{-1}$ (a) and soybean yields projected in corresponding sites (2000-2023) (b).**

Maize-soybean intercropping was sequentially allocated in highest yielding sites, ranked by soybean productivity based on mean average production over 2000-2023 period in rainfed conditions, until 50% soybean self-sufficiency (i.e., 18.1 Mt) is reached. All simulations are based on (i) yield projections from a random-forest model based on the two first principal components of climate variables and irrigation fraction, (ii) crop return frequency of one-in-four years, and (iii) partial land equivalent ratios equal to 0.56 and 0.79 for soybean and maize, respectively (12). Base map based on Natural Earth data, created using the R package rnaturalearth.

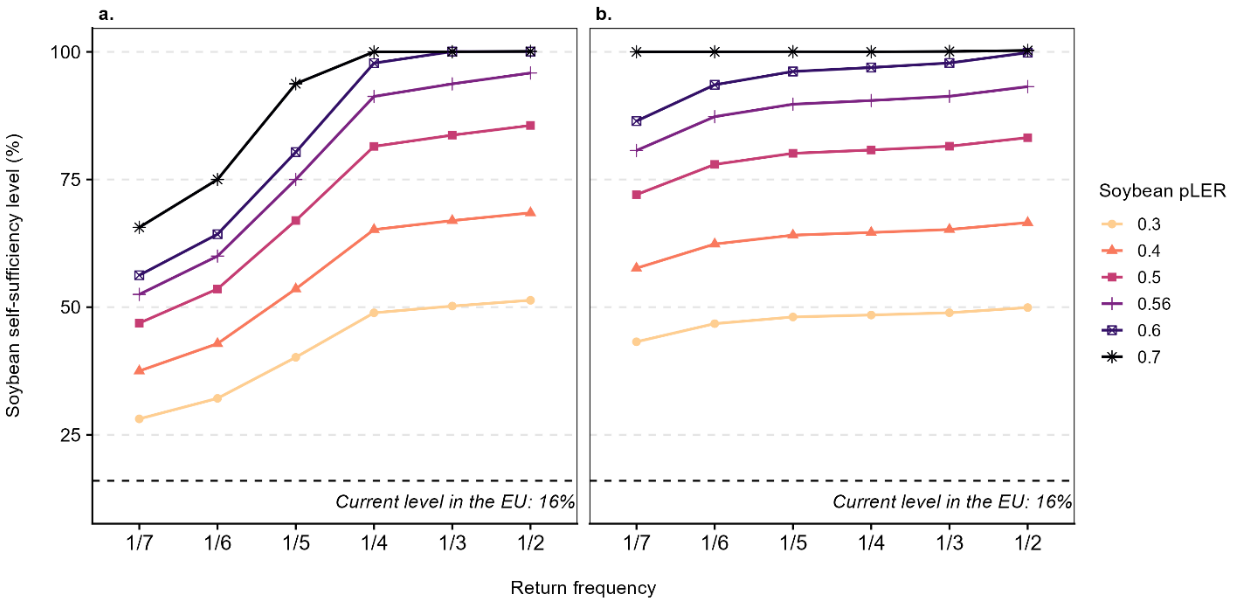


**Supplementary Figure 19. Level of soybean self-sufficiency in the European Union (EU) achieved from intercropping on areas showing soybean productivity equal or higher than 1 t.ha$^{-1}$ in the EU exclusively (left panel) or extended to neighboring countries (right panel), in several scenarios of partial land equivalent ratio (pLER) and crop return frequency.**

Crop frequencies of one year in seven, six, five, four, three, or two correspond to allocating maize-soybean intercropping on 14, 16, 20, 25, 33, or 50% of cropland area in each site. For all scenarios, total cropping area did not exceed 25% of croplands in the EU (i.e., 25 Mha). All simulations are based on (i) yield projections from a random-forest model based on the two first principal components of climate variables and irrigation fraction, (ii) crop return frequency of one-in-four years, and (iii) partial land equivalent ratios equal to 0.56 and 0.79 for soybean and maize, respectively (12).

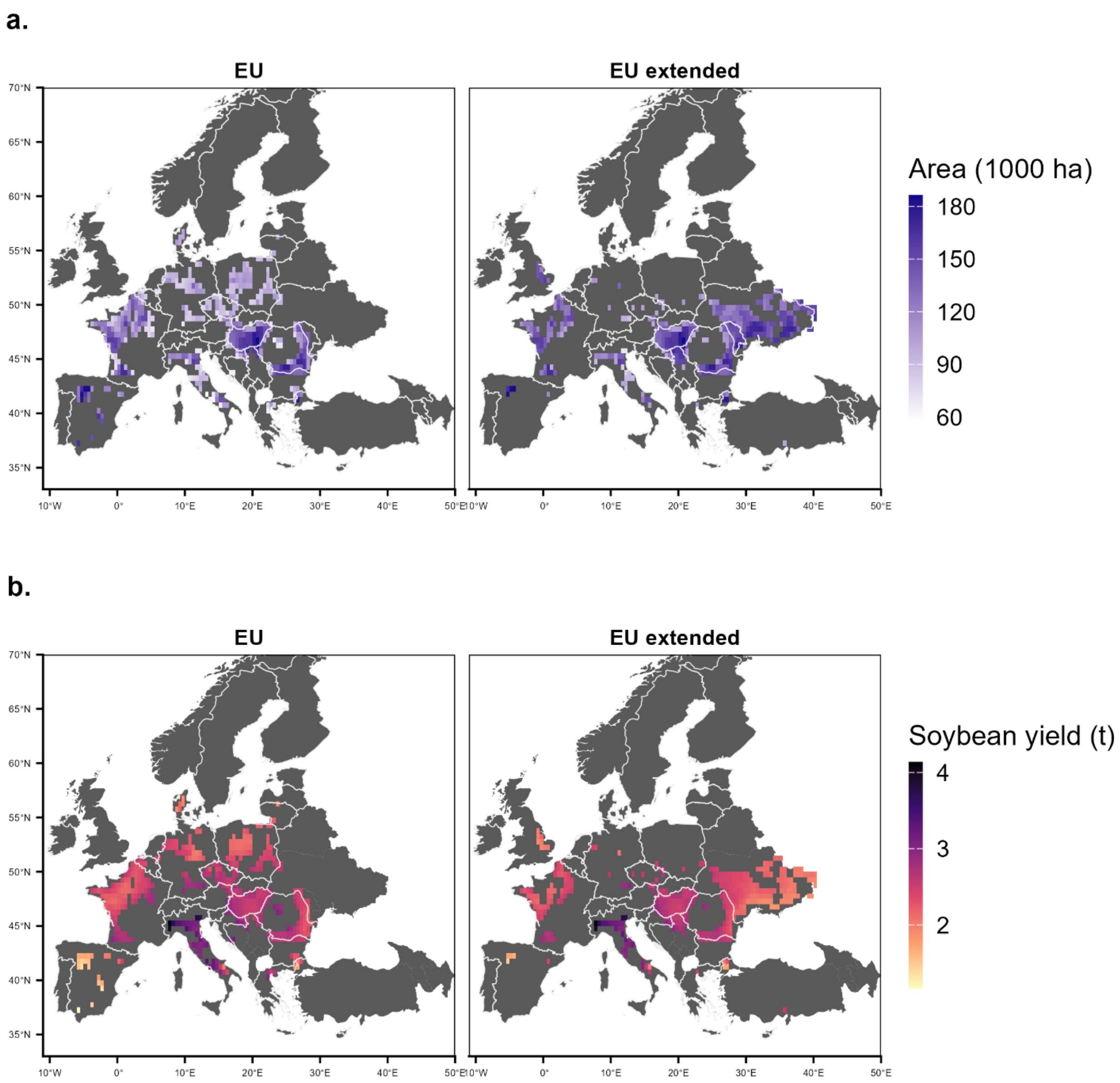


**Supplementary Figure 20. Geographical allocation of maize-soybean intercropping in the European Union exclusively (EU) or extended to neighboring countries (EU extended) (a) and soybean yields projected in corresponding sites (2000-2023) (b).**

Maize-soybean intercropping was sequentially allocated in highest yielding sites, ranked by soybean productivity based on mean average production over 2000-2023 period in rainfed conditions, until 100% soybean self-sufficiency (i.e., 36.3 Mt) is reached or 25% of EU' cropland area (i.e., 25 Mha) is covered. All simulations are based on (i) yield projections from a random-forest model based on the two first principal components of climate variables and irrigation fraction, (ii) crop return frequency of one-in-four years, and (iii) partial land equivalent ratios equal to 0.56 and 0.79 for soybean and maize, respectively (12). EU extended set of countries correspond to EU's members states and Azerbaijan, Belarus, Norway, Switzerland, United-Kingdom, Albania, Bosnia and Herzegovina, Georgia, Moldova, Montenegro, North Macedonia, Serbia, Türkiye, Ukraine, and Kosovo. Base map based on Natural Earth data, created using the R package rnaturalearth.

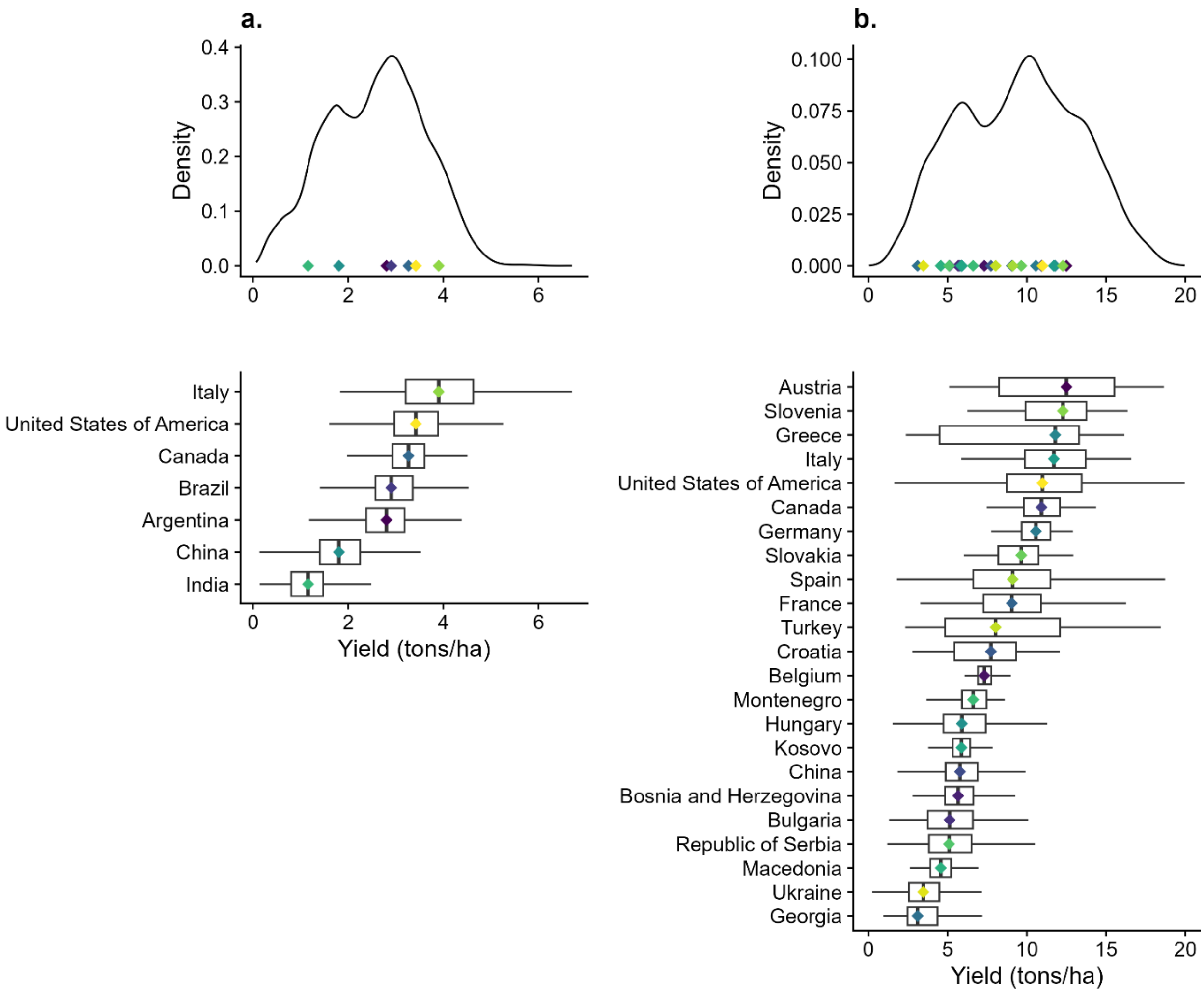


**Supplementary Figure 21. Soybean (a) and maize (b) yields distribution from 1981 to 2016 in sites located in producing areas included in the training dataset.**

Top panel: yield distribution is displayed as density for all sites located in producing areas for each crop; colored dots represent the median yield in each country included in the training dataset (excluding those located in unsuitable areas).

Bottom panel: yield distribution is displayed by country included in the training dataset (excluding those located in unsuitable areas) and ordered according to median yield.

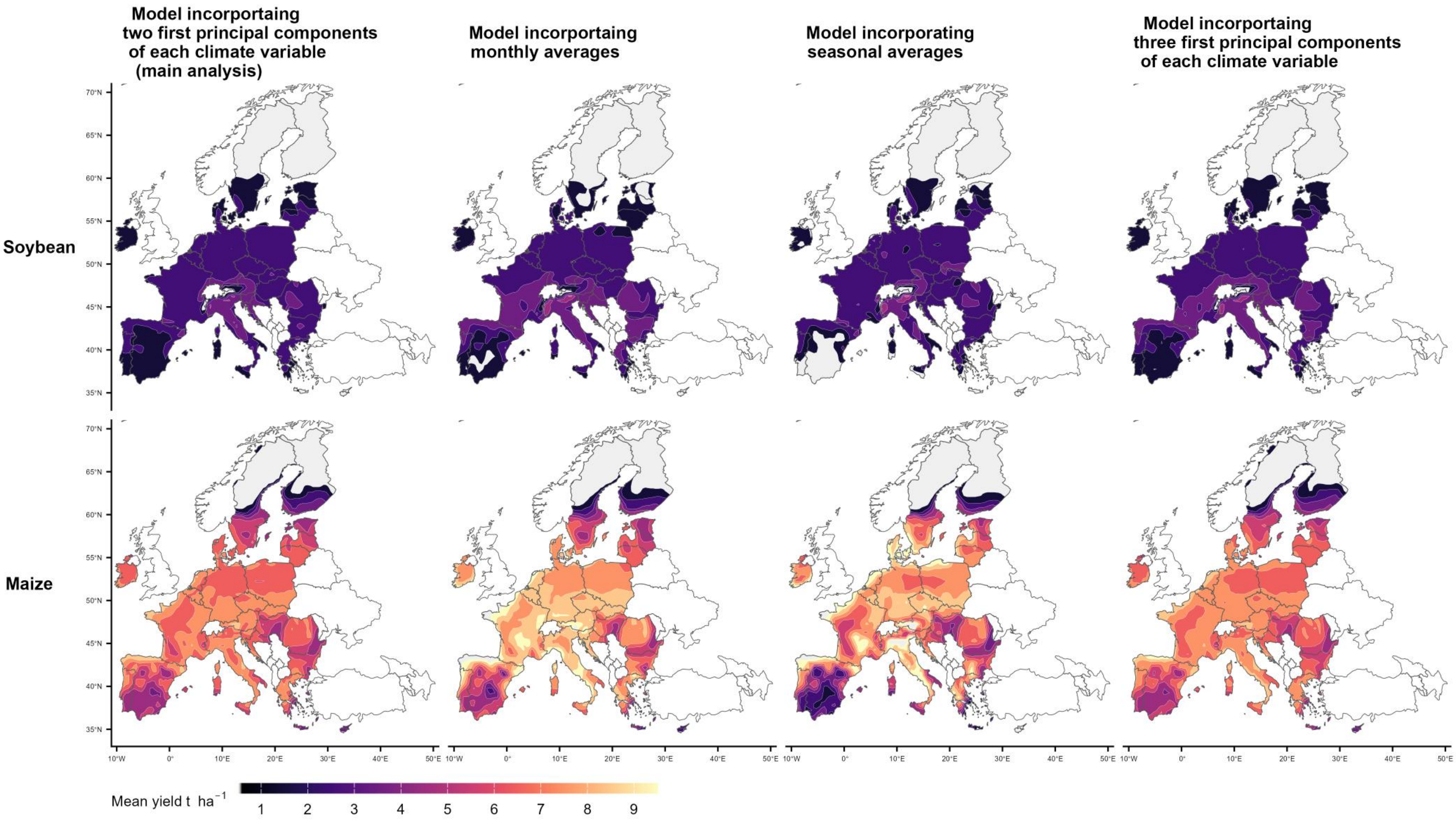


**Supplementary Figure 22. Projected yields of soybean and maize in the European Union according to four different predictive models.** Maps show average projected yields over 2000-2023 period in rainfed conditions. Projections were obtained using four random forest models taking into account climate inputs (derived from the ERA5-land dataset) covering the growing season (April to November for soybean and from April to December for maize) and irrigated fraction (estimated by the Spatial Production Allocation Model). Climate variables included minimum and maximum temperatures, total precipitations, solar radiation, vapor pressure deficit, evapotranspiration. Base map based on Natural Earth data, created using the R package rnaturalearth.

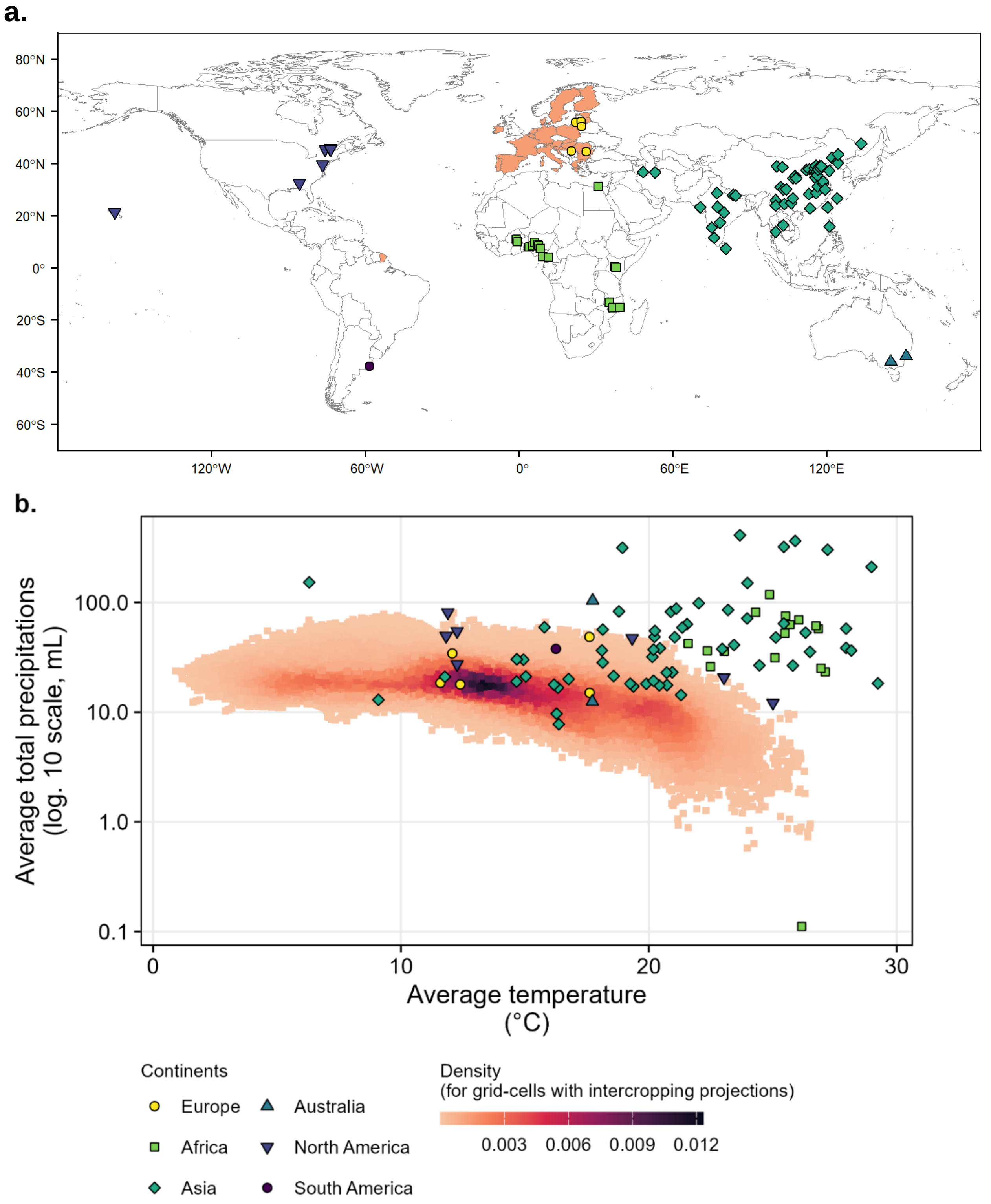


**Supplementary Figure 23. Location (a), average temperature and total precipitations during soybean growing period over 2000-2023 (b) in the experimental sites used in the meta-analysis of Xu et al. (12) and in the sites where soybean yields are projected in the present study.**
Cropping calendars for each site was determined using the USDA's crop calendars charts. For each site, monthly temperatures and total precipitations were derived from the ERA5 land dataset and then averaged over soybean growing period. Base map based on Natural Earth data, created using the R package rnaturalearth.

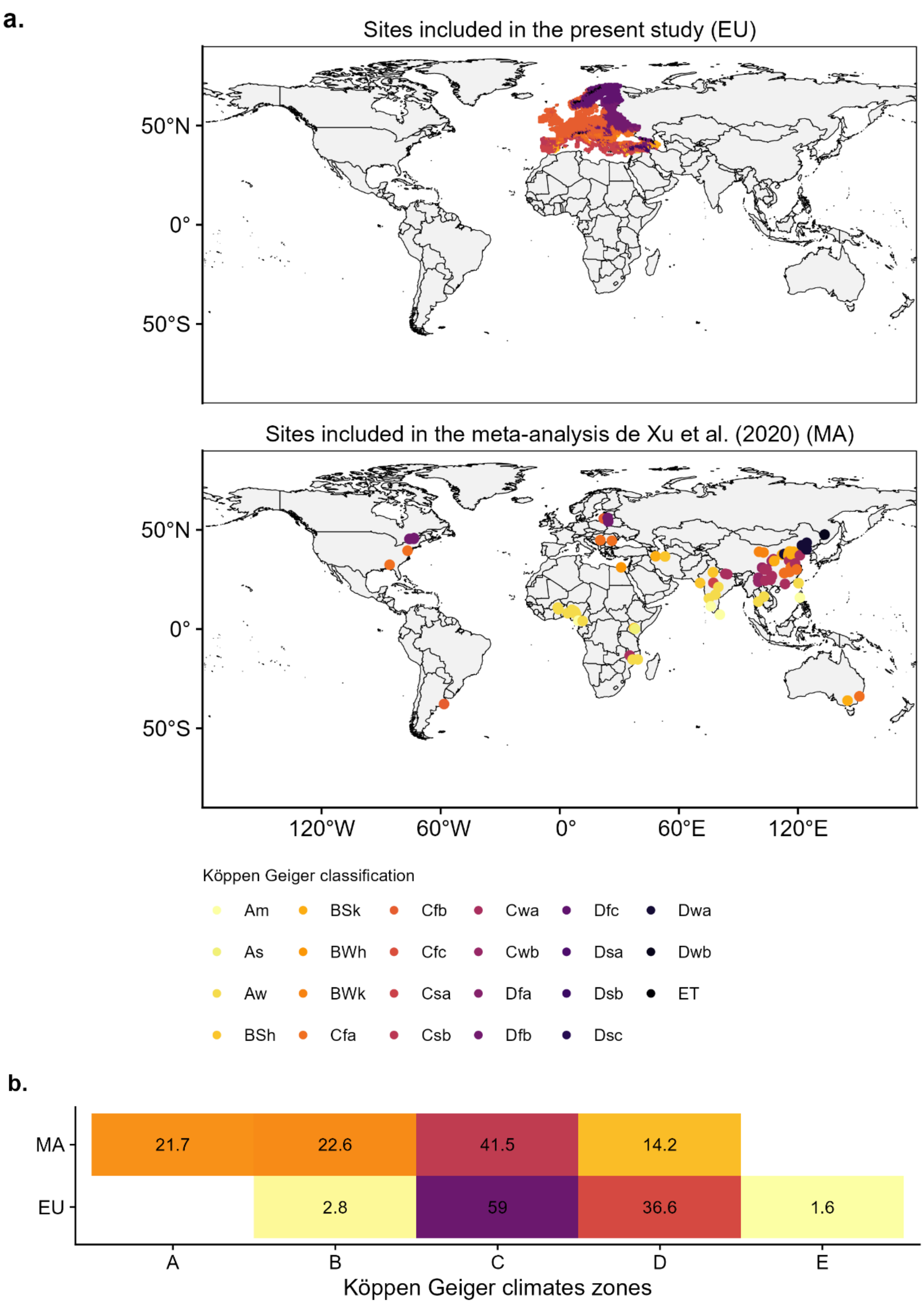


**Supplementary Figure 24. Location (a) and distribution (b) of Köppen-Geiger zones among the sites included in the meta-analysis of Xu et al. (12) [MA] and in the sites where soybean yields are projected in the present study [EU].** The numbers in (b) are the percentages of sites of the meta-analysis (MA) and in the projected European yield map (EU) located in each Köppen-Geiger zone. Base map based on Natural Earth data, created using the R package rnaturalearth.

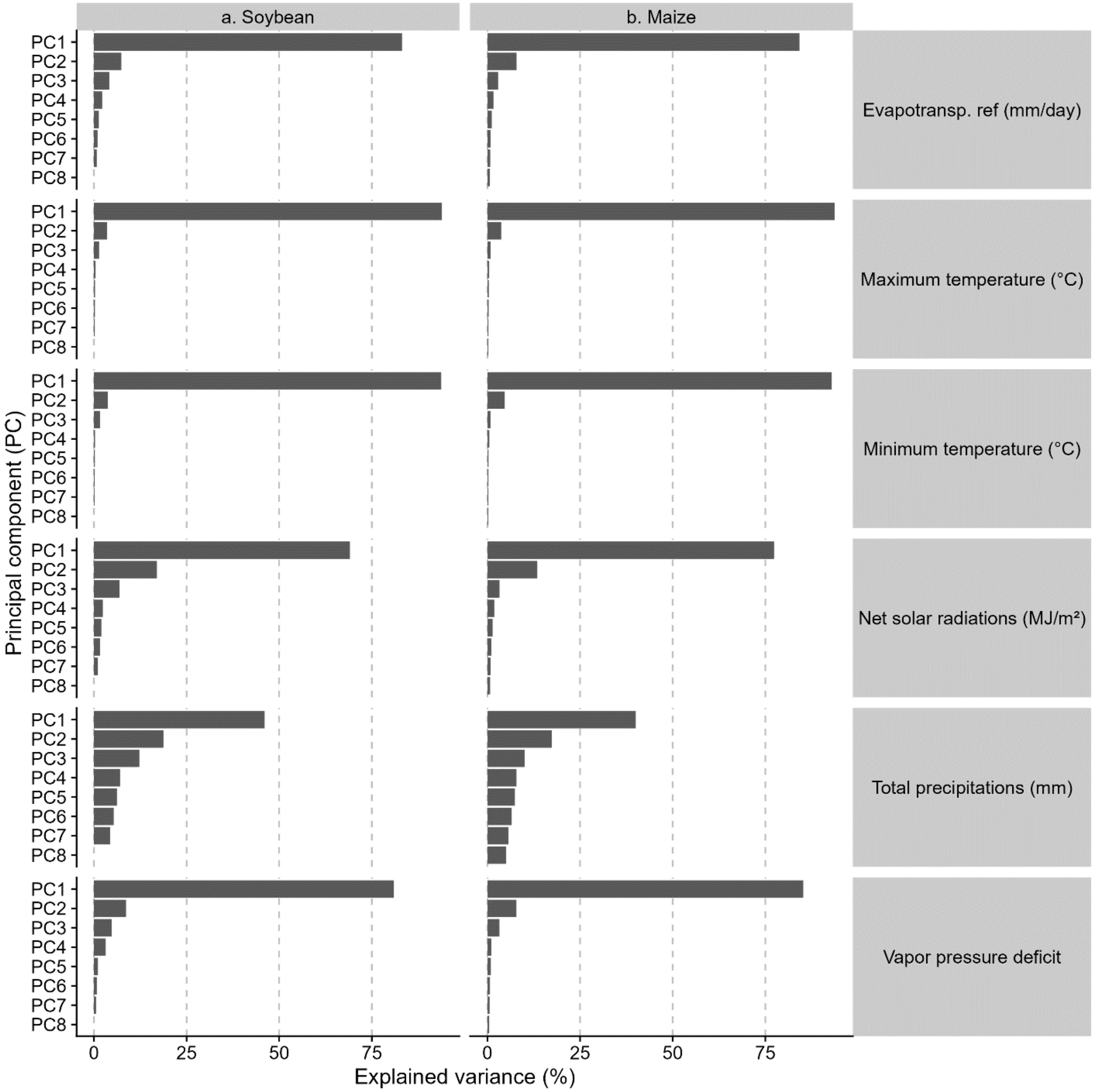


**Supplementary Figure 25. Variance explained by the principal components derived from principal component analysis applied on each climate variable in the soybean (a) and maize (b) training dataset.**

**Supplementary Table 1. Nash-Sutcliffe efficiency of climate-based models predicting soybean and maize yields estimated by cross-validation on years, sites, and on average.**

| | | Nash-Sutcliffe efficiency [c] computed by | | |
|---|---|---|---|---|
| **Climate-based crop yield predictive model [a]** | **Number of predictors [b]** | **cross-validation on years** | **cross-validation on sites** | **mean over both procedures** |
| **Soybean** | | | | |
| using the three first principal components derived from PCA applied on monthly climate data | 19 | 0.910 | 0.939 | 0.925 |
| ***using the two first principal components derived from PCA applied on monthly climate data*** | ***13*** | ***0.919*** | ***0.932*** | ***0.925 **** |
| using monthly climate data | 43 | 0.900 | 0.942 | 0.921 |
| using seasonal[d] climate data | 7 | 0.913 | 0.900 | 0.906 |
| **Maize** | | | | |
| using the three first principal components derived from PCA applied on monthly climate data | 19 | 0.902 | 0.911 | 0.907 |
| ***using the two first principal components derived from PCA applied on monthly climate data*** | ***13*** | ***0.911*** | ***0.905*** | ***0.908 **** |
| using monthly climate data | 49 | 0.866 | 0.917 | 0.892 |
| using seasonal[4] climate data | 7 | 0.920 | 0.872 | 0.896 |

Notes: climate data including minimum and maximum temperatures (both in °C), total precipitation (in mm), solar radiation (in MJ/m²), reference evapotranspiration (in mm/day), and vapor pressure deficit (in kPA). For each crop, * indicates the best predictive model based on average performance and number of predictors. Abbreviations: PCA: principal component analysis;

[a] All models use random forest algorithm.

[b] Including climate predictors and fractional area of irrigated crop.

[c] Ranging from 0 to 1, with higher value corresponds to better performance (an efficiency of 1 indicates perfect match between observations and predictions).

[d] Seasonal average refers to the mean over the growing season of the crop.

**Supplementary Table 2. Soybean and maize food balance, self-sufficiency rate, areas, and productivity in the European Union.**

| | Soybean [a] | Maize [b] |
|---|---|---|
| **Domestic supply breakdown by source,[c] in Mt of grains** | | |
| Production quantity | 2.7 | 66.5 |
| Export quantity | 11.6 | 25.0 |
| Import quantity | 45.2 | 44.8 |
| Stock variation | 0 | 1.1 |
| **Domestic supply quantity,[d] in Mt of grains** | **36.3** | **85.1** |
| **Mean yield,[e] in t ha$^{-1}$** | **2.8** | **7.5** |
| **Area harvested,[e] in Mha** | **1.0** | **8.9** |
| **Self-sufficiency rate in 2021-2022,[f] in %** | **16** | **81** |

[a] Including grains and cakes. Soybean cake quantity is expressed in soybean grain equivalent assuming that 1 Mt of soybean grain gives 0.8 Mt of soybean cake.

[b] Including maize, germ, flour, bran, gluten, starch, feed and meal, gluten (see definition and standards for the Food Balances domain).

[c] averages 2018-2022; data extracted from FAOSTAT: data extracted from “Food balances > Food balances (2010-)” domain (https://www.fao.org/faostat/en/#data/FBS).

[d] Computed as: Production quantity + Imports quantity + Stocks variation – Export quantity.

[e] averages 2018-2022; data extracted from FAOSTAT: data extracted from “Production > Crop and livestock products” domain (https://www.fao.org/faostat/en/#data/QCL).

[f] As reported in the first report on the State of Food Security in the EU of the European Food Security Crisis preparedness and response Mechanism (EFSCM) ‘A qualitative assessment of food supply and food security within the framework of the EFSCM’, available at: https://agriculture.ec.europa.eu/system/files/2023-11/efscm-assessment-autumn-2023_en.pdf

**Supplementary Table 3. Area of major crops in regions with high suitability of soybean cultivation in the European Union.**

| | Crop area in 2020 (in ha) in the regions with | | |
|---|---|---|---|
| | **projected soybean yield ≥1.0 $t.ha^{-1}$** | **projected soybean yield ≥2.1 $t.ha^{-1}$** | **projected soybean yield ≥2.6 $t.ha^{-1}$** |
| **Wheat** | 22.3 | 18.4 | 7.8 |
| **Maize (grain and forage)** | 11.8 | 11.0 | 6.5 |
| **Barley** | 10.5 | 6.7 | 2.7 |
| **Rapeseed** | 5.3 | 4.7 | 1.7 |
| **Sunflower** | 4.8 | 4.1 | 2.4 |

Notes: Data on crop area are taken from the CROPGRID gridded dataset (0.5°-resolution) (14). The soybean yield values of 1.0, 2.1, and 2.6 $t.ha^{-1}$ correspond to the $25^{th}$, $50^{th}$ (median), and $75^{th}$ percentiles of the rained soybean yield projections in the EU from a random-forest model based on the two first principal components of climate variables and irrigation fraction.

**Supplementary Table 4.** **Soybean and maize coproduction and surface requirement for 25, 50, 75, and 100% soybean self-sufficiency in the European Union (EU) achieved by intercropping or sole cropping *according to local nitrogen fertilization rates in the EU*.**

| | Target % of EU's self-sufficiency in soybean [a] | Intercropping | | Sole crops | | | Total | | Land-saving [b] |
|---|---|---|---|---|---|---|---|---|---|
| | | Soybean | Maize | Soybean | Maize | Additional land required | Intercrop-ping | Sole crop | |
| **Production (Mt)** | **25%** | 9.1 | 30.1 | 9.1 | 19.8 | 10.3 | 39.2 | 39.2 | |
| | **50%** | 18.2 | 67.3 | 18.2 | 44.7 | 22.6 | 85.5 | 85.5 | |
| | **75%** | 27.2 | 105.2 | 27.3 | 68.7 | 26.8 | 132.3 | 122.8 | |
| | **100%** | 31.0* | 121.9 | 36.3 | 65.0 | 5.4 | 152.9 | 106.7 | |
| **Surface (Mha)** | **25%** | 6.6 | | 3.4 | 3.2 | 1.5 | 6.6 | 8.1 | -19% |
| | **50%** | 13.9 | | 7.1 | 6.8 | 3.7 | 13.9 | 17.6 | -21% |
| | **75%** | 21.6 | | 10.9 | 10.7 | 4.2 | 21.6 | 25.9 | *-17%*** |
| | **100%** | 25.0 | | 14.8 | 10.2 | 0.9 | 25.0 | 25.9 | *-3%*** |

Notes: Crop return frequency was one-in-four year for all simulations, and total allocated area was limited to 25% of croplands in the EU (i.e., 25 Mha). For intercropping, partial land equivalent ratios were computed according to local nitrogen fertilization rates from the NPKGRID gridded dataset (11) based on the equation derived by Xu et al. (12). The computed values varied between 0.45 and 0.59 for soybean and between 0.77 and 0.79 for maize, respectively.

[a] Self-sufficiency levels of 50, 75, and 100% correspond to 18.2, 27.2, 36.3 Mt soybean.

[b] Computed as: ($surface_{\_intercropping}$.- $surface_{\_sole\ cropping}$)/ $surface_{\_sole\ cropping}$; representing the percentage of surface saved by intercropping compared to sole crop strategy.

* Target production not reached even when soybean was grown of 25% cropland in the EU and

** Production from intercropping strategy higher than sole crops.

**Supplementary Table 5. Soybean and maize coproduction and surface requirement for 25, 50, 75, and 100% soybean self-sufficiency in the European Union (EU) achieved by intercropping or sole cropping *assuming that maize and soybean are grown during the same period*.**

| | Target % of EU's self-sufficiency in soybean [a] | Intercropping | | Sole crops | | | Total | | Land-saving [b] |
|---|---|---|---|---|---|---|---|---|---|
| | | Soybean | Maize | Soybean | Maize | Additional land required | Intercrop-ping | Sole crop | |
| Production (Mt) | 25% | 9.1 | 29.6 | 9.1 | 20.4 | 9.3 | 38.7 | 38.8 | |
| | 50% | 18.2 | 65.5 | 18.2 | 45.1 | 20.4 | 83.6 | 83.7 | |
| | 75% | 27.2 | 101.7 | 27.3 | 68.5 | 27.0 | 128.9 | 122.8 | |
| | 100% | 31.1* | 118.1 | 36.3 | 65.0 | 5.3 | 149.2 | 106.7 | |
| Surface (Mha) | 25% | 6.8 | | 3.4 | 3.4 | 1.4 | 6.8 | 8.1 | -16% |
| | 50% | 14.1 | | 7.1 | 7.0 | 3.2 | 14.1 | 17.3 | -18% |
| | 75% | 21.6 | | 10.9 | 10.7 | 4.2 | 21.6 | 25.9 | *-17%*** |
| | 100% | 25.0 | | 14.8 | 10.2 | 0.9 | 25.0 | 25.9 | *-3%*** |

Notes: Crop return frequency was one-in-four year for all simulations, and total allocated area was limited to 25% of croplands in the EU (i.e., 25 Mha). Growing both soybean and maize during the same time period is translated by a temporal differentiation niche of 0. The values of partial land equivalent ratios of intercropping corresponding to this temporal differentiation niche were 0.53 and 0.76 for soybean and maize, respectively, based on the results of Xu et al. (12).

[a] Self-sufficiency levels of 50, 75, and 100% correspond to 18.2, 27.2, 36.3 Mt soybean.

[b] Computed as: (surface$_{\text{intercropping}}$.- surface$_{\text{sole cropping}}$)/ surface$_{\text{sole cropping}}$; representing the percentage of surface saved by intercropping compared to sole crop strategy.

* Target production not reached even when soybean was grown of 25% cropland in the EU.

** Production from intercropping strategy higher than sole crops.

*References*


1. Iizumi T & Sakai T (2020) The global dataset of historical yields for major crops 1981-2016. *Sci Data* 7(1):97.
2. Monfreda C, Ramankutty N, & Foley JA (2008) Farming the planet: 2. Geographic distribution of crop areas, yields, physiological types, and net primary production in the year 2000. *Global Biogeochemical Cycles* 22(1).
3. Chen M, Guilpart N, & Makowski D (2024) Comparison of methods to aggregate climate data to predict crop yield: an application to soybean. *Environmental Research Letters* 19(5):054049.
4. Guilpart N, Iizumi T, & Makowski D (2022) Data-driven projections suggest large opportunities to improve Europe's soybean self-sufficiency under climate change. *Nature Food* 3(4):255-265.
5. Dupin M*, et al.* (2011) Effects of the Training Dataset Characteristics on the Performance of Nine Species Distribution Models: Application to Diabrotica virgifera virgifera. *PLOS ONE* 6(6):e20957.
6. Kottek M, J. Grieser, C. Beck, B. Rudolf, and F. Rubel (2006) World Map of the Köppen-Geiger climate classification updated. . *Meteorol. Z.* 15:259-263.
7. Yu Q*, et al.* (2020) A cultivated planet in 2010 – Part 2: The global gridded agricultural-production maps. *Earth Syst. Sci. Data* 12(4):3545-3572.
8. Muñoz-Sabater J*, et al.* (2021) ERA5-Land: a state-of-the-art global reanalysis dataset for land applications. *Earth Syst. Sci. Data* 13(9):4349-4383.
9. Jolliffe I (2011) Principal Component Analysis. *International Encyclopedia of Statistical Science*, ed Lovric M (Springer Berlin Heidelberg, Berlin, Heidelberg), pp 1094-1096.
10. Breiman L. Random Forests. Machine Learning. 2001;45(1):5-32
11. Morales A, Villalobos FJ. Using machine learning for crop yield prediction in the past or the future. Frontiers in Plant Science. 2023;14.
12. Xu, Z. et al. Intercropping maize and soybean increases efficiency of land and fertilizer nitrogen use; A meta-analysis. Field Crops Research 246, 107661, doi:https://doi.org/10.1016/j.fcr.2019.107661 (2020).
13. Nguyen, T.H., Tang, F.H.M., Conchedda, G. et al. NPKGRIDS: a global georeferenced dataset of N, P2O5, and K2O fertilizer application rates for 173 crops. Sci Data 11, 1179 (2024). https://doi.org/10.1038/s41597-024-04030-4
14. Tang, X. *et al.* Intercropping legumes and cereals increases phosphorus use efficiency; a meta-analysis. *Plant and Soil* **460**, 89-104, doi:10.1007/s11104-020-04768-x (2021).
